# Hippocampal-amygdala interactions mediate uncertainty-dependent resistance to extinction following fear conditioning

**DOI:** 10.1101/725648

**Authors:** John Morris, Francois Windels, Pankaj Sah

## Abstract

The partial reinforcement extinction effect (PREE) is a paradoxical learning phenomenon in which omission of reinforcement during acquisition results in more persistent conditioned responding in extinction. Here, we report a significant PREE with an inverted-U, entropy-like distribution against reinforcement probability following tone foot shock fear conditioning in rats, which was associated with increased neural activity in hippocampus and amygdala as indexed by p-ERK and c-fos immunolabelling. *In vivo* electrophysiological recordings of local field potentials (LFPs) showed that 50% reinforcement was associated with increases in the frequency and power of tone-evoked theta oscillations in both the subiculum region of hippocampus and in basolateral amygdala (BLA) during both acquisition (Day 1) and extinction (Day 2) sessions. Tone-evoked LFPs in 50% reinforced animals also showed increases in coherence and bidirectional Granger Causality between hippocampus and amygdala. The results support a Bayesian interpretation of the PREE, in which the phenomenon is driven by increases in the entropy or uncertainty of stimulus contingencies, and indicate a crucial role for hippocampus in mediating this uncertainty-dependent effect.

## Introduction

The partial reinforcement extinction effect (PREE), in which omission of reinforcement during acquisition produces more prolonged conditioned responding in extinction, presents considerable problems for theories of associative learning (Rescorla and Wagner 1972) which propose that omission of reinforcement should weaken the association between conditioned stimulus (CS) and unconditioned stimulus (US) and thus result in more rapid extinction (Mackintosh, 1974, Lewis, 1960). Various alternative theoretical accounts of the PREE have been proposed, including frustration theory (Amsel 1958, 1992), sequential theory (Capaldi, 1966), rate estimation theory (Gallistel and Gibbon 2000) and Bayesian inferential theory (Courville et al., 2006). Frustration and sequential theories, while accounting for many features of the PREE, apply principally to appetitive instrumental conditioning protocols involving responses to rewards such as foods. These theories appear to have limited applicability to aversive classical conditioning protocols such as fear conditioning, where it is difficult to see how the omission of a foot-shock can give rise to an emotional state of “frustration”. Rate estimation and Bayesian inferential theoretical accounts of the PREE are both applicable to classical conditioning protocols but make distinct predictions about the presence and magnitude of the PREE.

Rate estimation theory proposes that, during conditioning, animals learn the temporal intervals between different events and the reciprocals of these intervals (i.e. rates) rather than form fixed stimulus-stimulus associations (Gallistel and Gibbon, 2000). Extinction occurs as the result of a decision process in which an animal stops responding after a fixed number of omissions of expected USs. If reinforcement is omitted on a proportion of acquisition trials (e.g. 20 CSs with only 10 USs), there will be a decreased reinforcement rate compared to continuous reinforcement (e.g. 20CSs with 20USs) and, the extinction “decision” will occur later than after continuous reinforcement. For example, in the above example, if an animal “decides” to extinguish after 10 expected USs, extinction will occur after 10 extinction CSs following continuous reinforcement, but after 20 CSs following the 50% partial schedule, i.e. a PREE will result. However, if US rates are made equivalent between partial and continuous protocols, rate estimation theory predicts a PREE will be absent.

Bayesian inferential theory proposes that subjects in a conditioning situation possess a “world model” whose parameters specify contingencies between stimuli and reinforcement, and which is continually updated in the light of new events. Significantly, updating of the world model depends on the level of uncertainty: in uncertain situations, more weight is given to novel events. In the context of the PREE, uncertainty will be greater during partial than continuous reinforcement in acquisition, and therefore unexpected US omissions will be given extra weight in updating the world model parameters, i.e. learning will be enhanced for partially reinforced subjects; in extinction, US omission in continually reinforced subjects will be surprising, evoking uncertainty that will enhance learning, i.e. accelerate extinction, whereas in partially reinforced subjects, US omission will not be surprising and extinction learning will be slower. Uncertainty, as measured by Shannon entropy (H(x) = -∑p(x) log(p(x)), where p is the probability mass function of x (Shannon, 1948), describes an inverted U function across probability with a peak at 0.5. Bayesian inferential theory, therefore, predicts that the PREE should vary in magnitude across partial reinforcement schedules in an inverted-U fashion with a peak at 50%.

Previous research into the neural basis of the PREE has almost exclusively employed appetitive instrumental protocols (Rawlins et al. 1980; Rawlins et al. 1989, Jarrard et al. 1986; Gray et al., 1972; Gray, 1970; Gray et al., 1975). These studies indicate that the hippocampus, and in particular hippocampal theta oscillations, may play a critical role in mediating the effect (Rawlins et al. 1980; Rawlins et al. 1989, Jarrard et al. 1986; Gray et al., 1972; Gray, 1970). Lesions of the hippocampus, and its main output structure, the subiculum, abolish the PREE (Rawlins et al. 1980; Rawlins et al. 1989, Jarrard et al. 1986). Induction of theta oscillations in hippocampus, via low frequency electrical stimulation of the septum, produces the equivalent of a PREE (Gray, 1970), whereas high-frequency stimulation of the septum, which blocks hippocampal theta, abolishes the PREE (Gray et al., 1972; Gray, 1970). Administration of the barbiturate, amobarbitol, which increases the threshold for septal driving of hippocampal theta, also abolishes the PREE (Gray, 1970). It is unknown whether hippocampus and hippocampal theta oscillations are also crucial for the expression of a PREE resulting from aversive classical conditioning (e.g. fear conditioning) and also whether PREE-related theta is involved in synchronizing activity across separate brain regions, as has been shown for other learning phenomena (Narayanan et al., 2007; Seidenbecher et al., 2003).

In the first experiment of the current study, rats were fear conditioned using both partial (25%, 50% and 70%) and continuous (100%) reinforcement schedules and neural activity during extinction was measured using immunohistochemical labelling for p-ERK and c-fos. In the second experiment, *in vivo* electrophysiological recordings were made during both acquisition and extinction in the hippocampus and amygdala of rats undergoing 50% and 100% reinforcement schedules. These experiments were designed to determine: firstly, whether a PREE can be demonstrated in a standard tone foot shock fear conditioning protocol in rats; secondly, whether rate estimation theory or Bayesian inferential theory provide a better theoretical account of the phenomenon; thirdly, how neural activity in amygdala and hippocampus, and in particular theta oscillations, correlate with the PREE; and finally, whether interactions between amygdala and hippocampal neural responses, as indexed by coherence and Granger Causality analyses, are associated with the PREE.

## Methods: Experiment 1

### Subjects

Twelve 10 week old adult male Wistar rats (weight 330-400g) were housed under standard laboratory conditions with a 12 hour light/dark cycle and food and water freely available. All experimental procedures were approved by the research ethics committee at the University of Queensland.

### Materials

All conditioning was conducted in MedAssociates (St Albans, VT, United States) conditioning chambers (30×24×21cm), and subjects’ behaviour was recorded with digital video cameras. A 20 second 80dB 5000Hz tone was used as the conditioned stimulus (CS) and a 0.5s 0.6mA foot shock as the unconditioned stimulus (US). Following conditioning, immunohistochemical labelling was undertaken using primary antibodies for c-fos (1:500, Cell Signaling) and p42/44 MAPK (Erk1/2) (1:500, Cell Signaling).

### Behavioural protocol

Subjects were divided into four experimental groups (three per group), each of which received a different fixed percentage (either 25%, 50%, 70% or 100%) of reinforcement during acquisition of conditioning. However, the absolute number and timing of reinforced trials were made identical across all groups, i.e. 7 CS-US pairings. In order to achieve the different percentages of reinforcement, the number of unreinforced trials (i.e. CS alone) during the session was varied across groups. Thus, while the 100% group experienced a total of 7 CS presentations in the acquisition session (all paired with the US) the 70% group received an additional 3 unreinforced CS presentations (total 10 trials), the 50% group an additional 7 unreinforced CS presentations (total 14 trials) and the 25% group an additional 21 unreinforced CS presentations (total 28 trials). The session length (90 minutes) was the same for all groups, and therefore, in addition to a variation in the total number of trials between groups, the inter-trial interval also varied: 2-4 minutes for the 25% group, 4-12 minutes for 50%, 4-18 minutes for 70%, and 4-18 minutes for 100%. The extinction protocol was the same for all groups, consisting of 28 presentations of the CS tone without the shock US over the course of a 90 minute session (inter-trial interval 2-4 minutes). Freezing (Bolles 1970) and crouching (Blanchard and Blanchard 1969; Blanchard and Blanchard 1972) behaviour were used to measure conditioned responding (CR). Freezing (crouching) was assessed by manual observation on a second by second basis carried out by an experimenter blind to subject group.

### Immunohistochemistry

Two hours after the extinction session, subjects were anaesthetized with isofluorane and given an intraperitoneal injection of Lethobarb (0.5 – 1.0ml). They were then perfused pericardially with 4% paraformaldehyde in 0.1M phosphate buffer and their brains removed and post-fixed overnight. After immersion in sucrose in phosphate buffer, the brains were frozen with liquid nitrogen and coronally sectioned (50μm) with a vibrotome (Leica). A sample of 1 in 6 sections was then processed for immunohistochemistry. After rinsing with phosphate buffered saline (PBS) and blocking with 3% bovine serum albumin, sections were incubated with primary antibodies for c-fos. After washing in PBS, sections were incubated in biotinylated secondary goat anti-rabbit antibody. ABC peroxidase complex (Vector) was used for signal amplification and after washing in PBS, the peroxidase end product was visualized with DAB (0.025%) and 5% nickel ammonium sulphate. The sections were then secondarily labelled with p42/44 MAPK (Erk1/2) antibody. The previous steps were repeated for the c-fos labelling with the substitution of c-fos for the p42/44 MAPK (Erk1/2) antibody. Finally, the sections were mounted on gelatin-coated slides and coverslipped. The sections were scanned and labelled cell densities measured using unbiased stereological techniques. Regions of interest were identified according to the rat atlas of Paxinos and Watson (2006). Counting of c-fos and p-ERK labelled cells was performed by an experimenter blind to the experimental group. C-fos positive cells were identified by intense, black nuclear staining, while p-ERK positive cells were identified by a more uniform brown staining of both cytosol and nucleus. All immunopositive cells within each region of interest on every section were counted, i.e. there was no sampling of areas within regions of interest. The effect of changes in probability of reinforcement on freezing behaviour and immunolabelling was tested with one-way analysis of variance (ANOVA). In order to test the statistical significance of differences between specific groups, comparisons were made using Student’s t test.

## Methods: Experiment 2

### Subjects

Twelve adult male Wistar rats, 8-12 weeks of age (350-420g) were housed under standard laboratory conditions with a 12 hour light/dark cycle and food and water freely available. Three subjects died following surgery and before the start of conditioning acquisition, leaving nine subjects from which behavioural and electrophysiological data were collected. All experimental procedures were approved by the research ethics committee at the University of Queensland.

### Materials

Nickel-chromium microelectrode wire with a 12.5 micron diameter core was used to assemble tetrodes (twisted bundles of four wires) using a custom-made apparatus. Eight tetrodes were then assembled into a micro-drive array. Prior to surgery, the electrodes were plated with gold and impedances measured. Conditioning was undertaken in a circular arena (diameter 75cm) with a metal plate floor and a Perspex wall.

### Surgery

Subjects underwent surgery for implantation of the microelectrode array about two weeks prior to the acquisition of conditioning. After a suitable level of anaesthesia under urethane was obtained, subjects were placed in stereotaxic frame and the skull exposed. Holes were drilled in the skull at coordinates corresponding to the brain regions of interest for that particular subject. The head stage was securely fixed with dental cement at the end of surgery.

The placement of tetrodes in each of the subjects was as follows: Subject 1: 4 tetrodes only, all in BLA; Subject 2: 4 tetrodes in BLA, 4 in subiculum; Subject 3: 4 tetrodes in BLA, 4 in central amygdala; Subject 4: 4 tetrodes in BLA, 4 in central amygdala; Subject 5: 4 tetrodes in BLA, 4 in central amygdala; Subject 6: 4 tetrodes in BLA, 4 in subiculum; Subject 7: 4 tetrodes in BLA, 4 in subiculum; Subject 8: 4 tetrodes in BLA, 4 in subiculum; Subject 9: 4 tetrodes in BLA, 4 in subiculum; Only data from the BLA and subiculum electrodes are reported here.

Following post-operative recovery, the subjects were handled regularly by the experimenters and the microelectrodes advanced on a daily basis. Several days prior to the acquisition of conditioning, subjects were placed in the experimental conditioning apparatus, connected via a cable to the recording equipment and brief electrophysiological measurements were made to assess whether the microelectrodes had reached an appropriate depth for the conditioning experiment to start. No tones, electric shocks or other stimuli were presented during these brief screening sessions.

### Histology

After the experiment was completed, the subjects were sacrificed, perfused pericardially with 4% paraformaldehyde in 0.1M phosphate buffer and their brains removed and post-fixed overnight. After immersion in sucrose in phosphate buffer, the brains were frozen with liquid nitrogen and coronally sectioned (50μm) with a vibrotome (Leica). Sections were examined with microscopy in order to determine the anatomical location of the tetrodes.

### Behavioural protocol

Once a subject had been sufficiently habituated to the experimenters and the experimental apparatus, and it was determined from the screening sessions that the microelectrodes were optimally placed, acquisition of conditioning commenced. The 50% and 100% behavioural protocols were similar to Experiment 1 except that there were 27 rather than 28 trials in extinction in the present experiment. Five subjects followed the 50% protocol (Subjects 2,3,4,10,12; four subjects followed the 100% protocol (Subjects 5,6,7,8).

The standard 100% acquisition protocol consisted of seven 5000Hz 80 dB pure tones each of 20 second duration (CS), and co-terminating in a 0.5 second 0.6mA foot shock (US), all delivered during the course of a 90-minute session. Due to a technical problem, the seventh trial was omitted in two of the subjects (subjects 6 and 7), giving them six rather than seven tone-shock pairs in acquisition. The data from individual subjects were analyzed separately and compared. No significant differences were evident between the two subjects with seven trials and the two subjects with six trials, and therefore the data from all four subjects were combined in the 100% group.

The standard 50% acquisition protocol consisted of 14 5000Hz 80dB pure tones of 20 second duration (CS), seven of which coterminated in a 0.5 second 0.6mA foot shock (US), delivered during the 90 minute session. The timing of the seven reinforced trials coincided with those in the 100% condition. Due to technical problems, a fifteenth unreinforced tone was presented at the end of the acquisition session in two subjects (subjects 8 and 9). Comparison of individual subject data between the three subjects receiving 14 and the two receiving 15 trials revealed no significant differences, and therefore the data from all subjects were combined in the 50% group. In subject 10, there was a technical problem with the timing of some of the shocks during reinforced trials, so that they were delivered a few seconds early. In the case of subject 10, therefore, the tone-evoked activity in the affected trials was measured over a shorter epoch (the shortest epoch being 13.5 seconds) than the standard 19.5 seconds. No significant differences were evident between the individual data of subject 8 and the other 50% condition subjects, and the data from this subject were combined with the 50% group.

The extinction protocol was the same for both 50% and 100% conditions, consisting of 27 presentations of the CS tone without the shock US over the course of a 90 minute session (inter-trial interval 2-4 minutes). In subjects 2 and 3, the extinction session consisted of a repeat of the 50% acquisition sequence with omission of the US. Consequently, subjects 2 and 3 received 14 unreinforced trials rather than the 27 received by subjects 3-9. A separate analysis of subjects 2 and 3 showed no significant difference from the results of the other 50% subjects and data from all subjects were combined in the 50% group.

### Electrophysiology

Local field potentials (LFP) were recorded on a subset of the 32 channels potentially available. The channels used for LFP were determined at the time of data acquisition, depending on how many channels were active, which channels were used for reference and which channels were chosen to record unit activity. Data were acquired using the Axona Dacq USB recording system. A preamplifier/analogue to digital converter, operating at a fixed gain of 100, digitized the signals from the microelectrodes with 24 bit resolution and a 48 kHz sampling rate. The LFP signals were routed to the system unit where they were low pass filtered at 300 Hz. The filtered signal was then stored as a .BIN file on the hard drive of a PC via a USB2 connection. The Axona system unit controlled the delivery of tone and shock stimuli and also the recording of the overhead behavioural tracker video.

### Data analysis

The low pass filtered raw signal in the .BIN file was imported into MATLAB using a custom written script providing 32 channels of data with a 48 kHz sampling rate for each subject (16 for subject 1). After removal of reference channels, channels reserved for unit recording data and channels with no signal, the remaining channels with usable LFP signals were submitted for data analysis. As a first step, the data were formatted for importation into the EEGLAB toolbox for MATLAB (http://sccn.ucsd.edu/eeglab/), and then resampled at a rate of 1500Hz. For a basic spectral analysis, a Fourier transform with a Hanning window of 6 seconds, a 2 second overlap, and a frequency range of 0.3 – 45 Hz was applied to the LFP data from both acquisition (Day 1) and extinction (Day 2). Epochs were extracted from these spectrograms corresponding to the 18 seconds of the tone CS preceding the shock US, and a pretone period also of 18 seconds.

Following this basic spectral analysis, a statistical comparison (ANOVA) of the 50% and 100% group LFPs was undertaken for the theta frequency range 8-10Hz in both acquisition and extinction sessions. As a first step, the power in the LFP theta band in each pretone and tone epoch was normalized by expressing it as a ratio of the mean spectral power in that epoch. These normalized data were then used to derive tone-evoked changes in LFP theta power for each trial by expressing the normalized theta power in each tone epoch as a ratio of the normalized theta power in the preceding pretone epoch.

An analysis of coherence between BLA and subiculum LFP channels was implemented using the EEGLAB toolbox for MATLAB. Coherence was first calculated in all subjects for pretone and tone epochs separately over a broad frequency spectrum (0.3 – 45 Hz) and in both acquisition and extinction sessions using the 1500Hz resampled LFP data. A statistical comparison (ANOVA) of 50% and 100% group LFP coherence was undertaken for the theta frequency range 8-10Hz. Coherence in the LFP theta band was first calculated for each pretone and tone epoch, and then the tone-evoked coherence calculated by expressing the tone coherence as a ratio of the pretone coherence.

A Granger Causality analysis was undertaken with a Granger Causality Open Source MATLAB toolbox (Luo, et al., 2013), using the same BLA and subiculum channels used in the coherence analysis. The original 48kHz LFP data were downsampled to 480Hz using the functionality of the toolbox. A partial Granger Causality analysis for each pretone and tone epoch was performed with an Autoregressive Integrated Moving Average (ARIMA) model order of 4. A statistical analysis (ANOVA) of the Granger Causality values for the 50% and 100% groups was undertaken for both acquisition and extinction sessions and in both BLA=>subiculum and subiculum=>BLA directions.

## Results: Experiment 1

### Behaviour

Freezing behaviour to the tone CS in trials 8-21 of the extinction session was significantly greater following 50% reinforcement than 100% reinforcement (p<0.05, t test, Figure 1). There was an overall significant main effect of reinforcement probability on freezing behaviour during trials 8-21 of the extinction session (ANOVA, F=4.338, p=0.0431, n=12, Figure 2). These behavioural findings, therefore, provide the first report of a significant PREE in tone foot-shock fear conditioning in rats. The plot of freezing behaviour against reinforcement probability formed an inverted U distribution (Figure 2).

**Figure 1.**
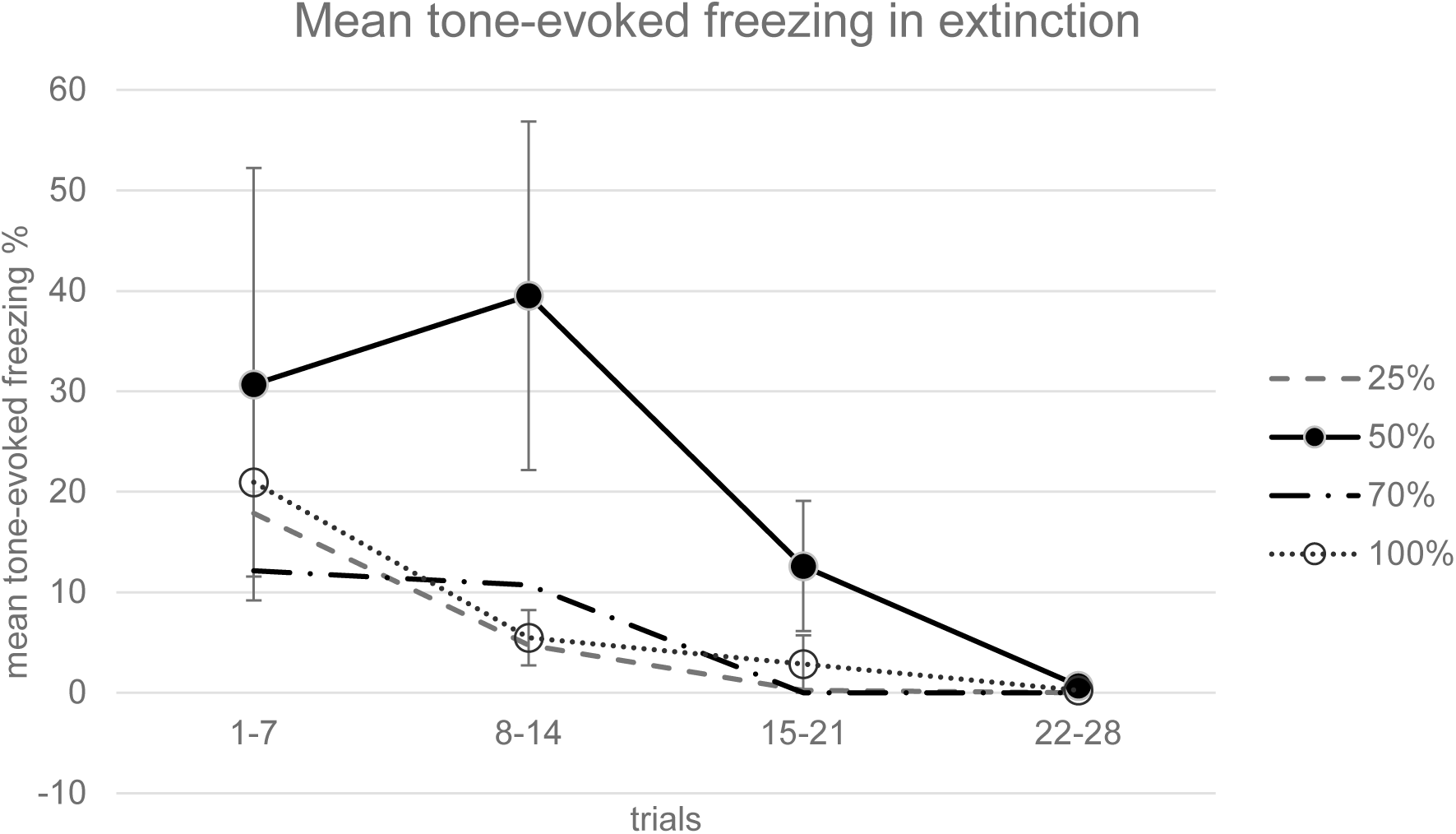
Timecourse of group tone-evoked freezing (crouching) during the extinction session (timecourse). Mean percentage of time spent freezing during the tone is displayed for the first trial and for 7 trial bins in each reinforcement group. Tone-evoked freezing in the 50% group was significantly greater (p<0.05) than that in the 100% group in trials 8-21 with a one-tailed Student’s t test; n=3 for each group. Error bars indicate +/−1 standard error.

**Figure 2.**
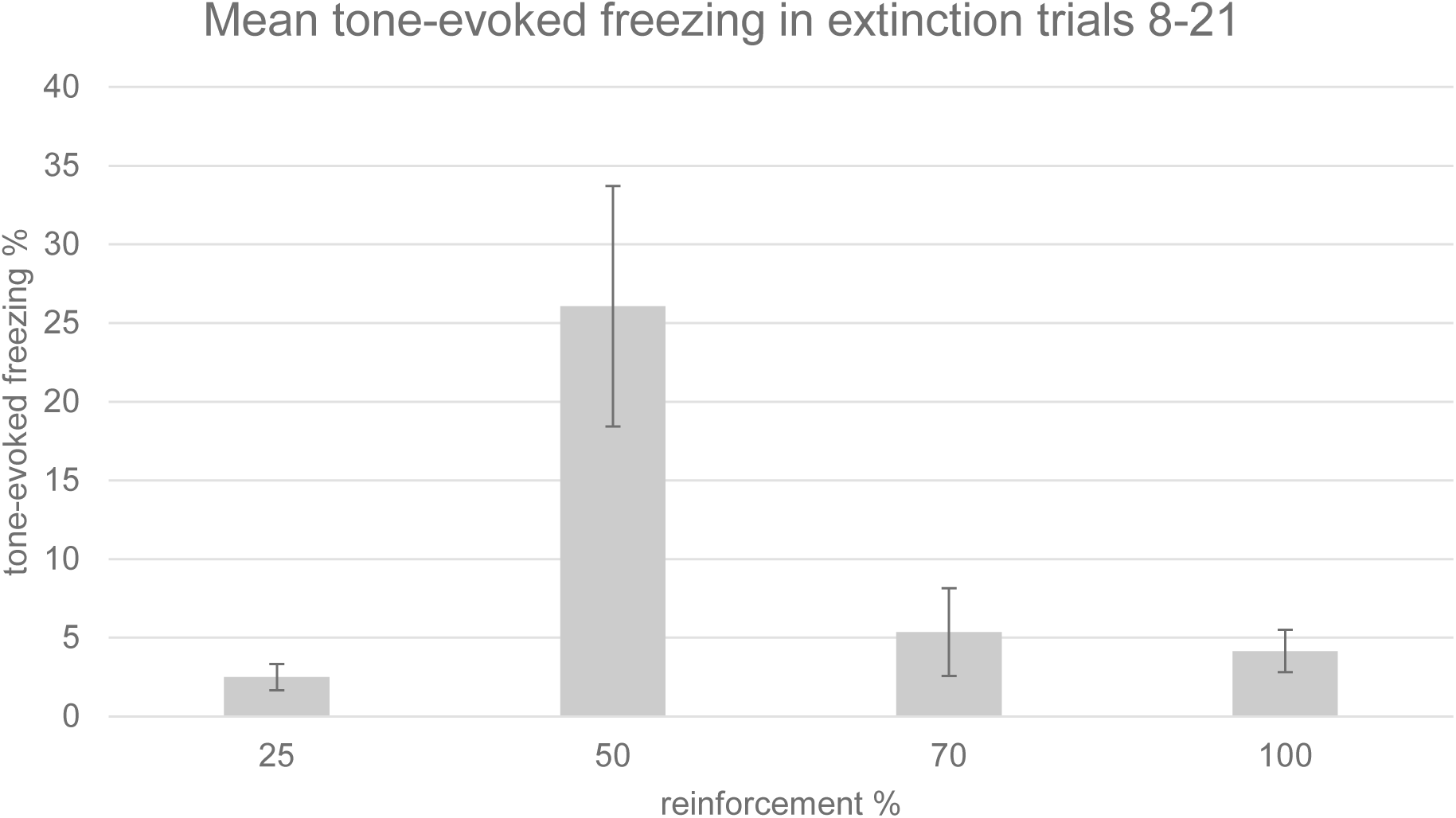
Group mean tone-evoked freezing (crouching) during the extinction session. Freezing was assessed manually by an experimenter blind to the experimental conditions. Mean percentage of time spent freezing is displayed for trials 8-21, in each reinforcement group. There was a significant effect of reinforcement probability on freezing (ANOVA, F=4.338, p=0.0431). Tone-evoked freezing in the 50% group was significantly greater (p<0.05) than that in the 100% group in trials 8-21 with a one-tailed t test; n=3 for each group. Error bars indicate +/−1 standard error.

### Immunohistochemistry

Representative coronal sections of the amygdala from subjects from each of the condition groups (25%, 50%, 70% and 100%) are shown in Figure 3. Prominent labelling of p-ERK and c-fos in the central nucleus of amygdala is evident in the 50% condition, with lesser degrees of labelling in the other conditions, particularly 100%. In the capsular subdivision of the central amygdala (ANOVA, p>0.05), p-ERK labelling was significantly greater for partial group subjects (25%, 50% and 70%) compared with the continuous (100%) group (t-test, p<0.05, Figure 4) although there was no significant main effect of reinforcement probability on p-ERK or c-fos labelling (ANOVA, p>0.05). Individual subject p-ERK labelling in capsular central amygdala in extinction correlated significantly with freezing behaviour (ANOVA, F=14.82, p<0.01, Figure 5). Labelling of p-ERK and c-fos in basolateral amygdala (BLA) showed no significant effects of reinforcement probability (ANOVA, p>0.05, Figure 6). While there was no significant main effect of reinforcement group on p-ERK or c-fos cell density in ventral hippocampus (ANOVA, p>0.05), p-ERK labelling in the 50% group was significantly greater than the 25%, 70% and 100% (t test, p<0.05, n=12, Figure 7). C-fos labelling in the CA1 region of ventral hippocampus was significantly correlated with c-fos labelling in BLA in the extinction session (ANOVA, F=5.31, p=0.05), Figure 8).

**Figure 3.**
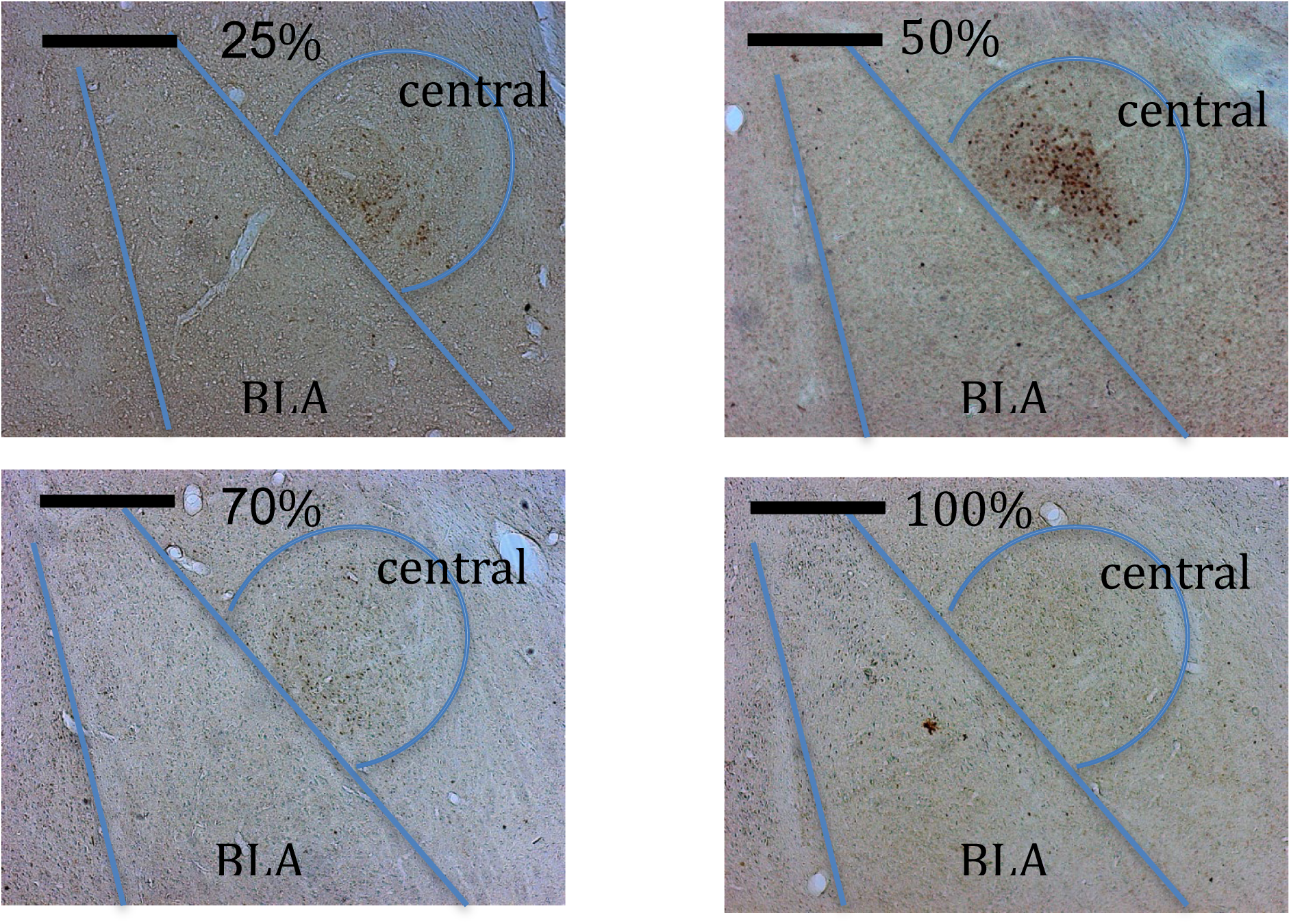
Representative 50μm coronal sections of amygdala stained for both c-fos and p-ERK. Sections from rats in all four reinforcement groups are shown: 25%, 50%, 70%, and 100%. Scale bar 500µm.

**Figure 4.**
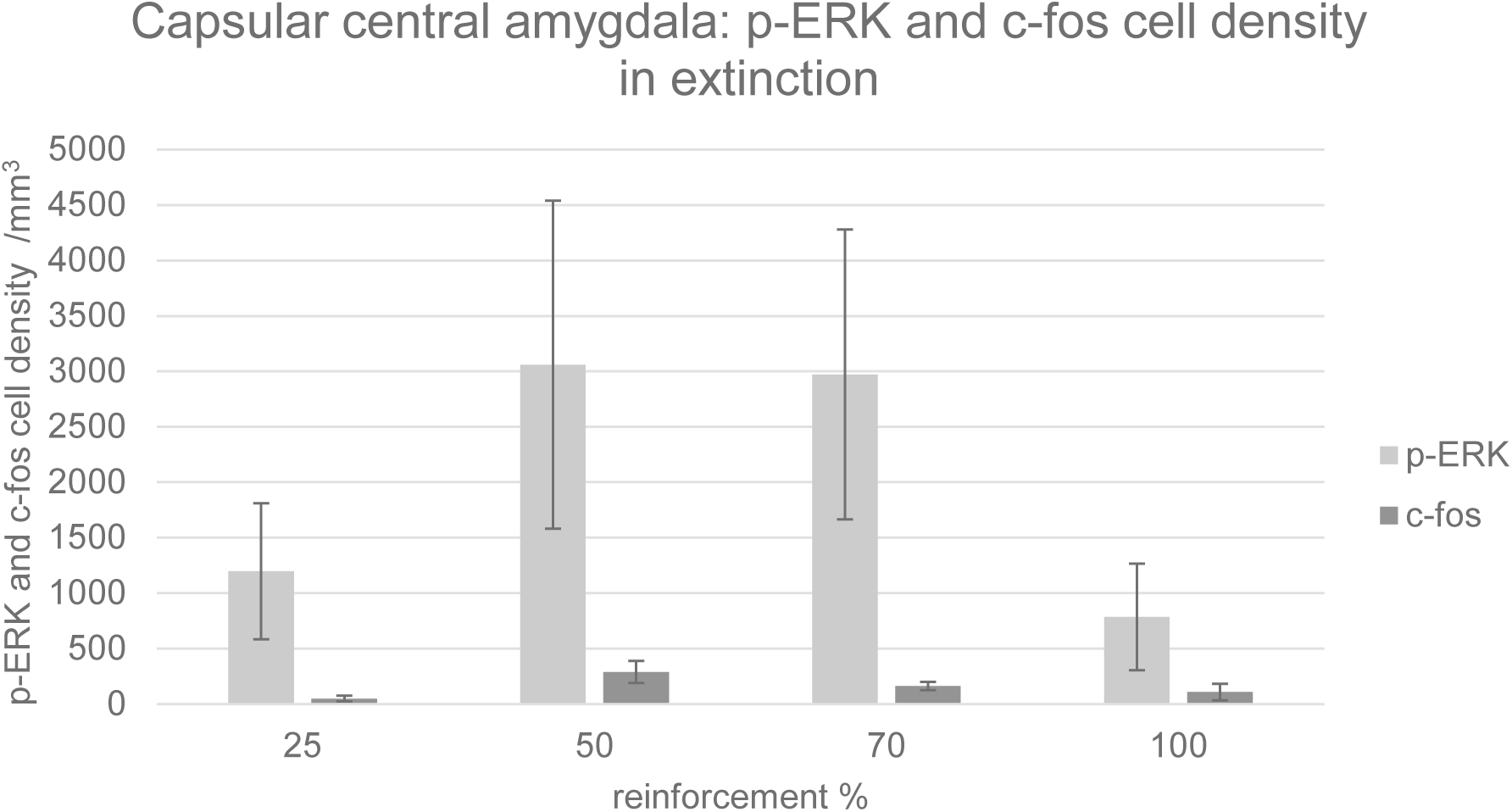
Density of p-ERK and c-fos labelled cells per mm^3^ in capsular subdivision of central amygdala. Error bars indicate +/−1 standard error. There was no significant main effect of reinforcement group on p-ERK or c-fos cell density (ANOVA, p>0.05). p-ERK labelling in the partially reinforceed groups (25%+50%+70%) was significantly greater than the 100% group in a one-tailed Student’s t test (p<0.05); n=3 for each group.

**Figure 5.**
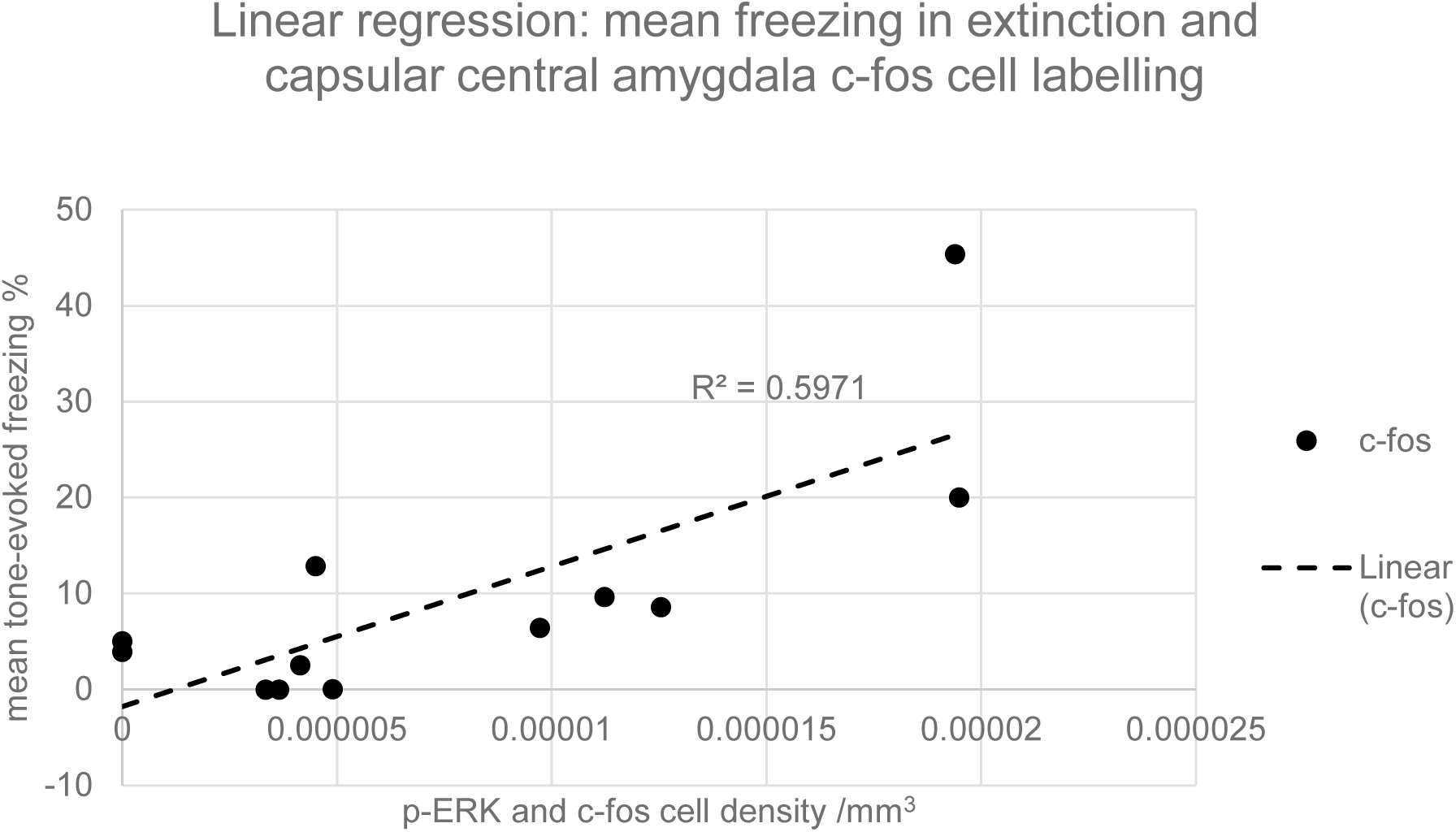
Graph of linear regression between mean c-fos cell density in capsular central amygdala and the mean tone-evoked freezing in extinction for all subjects. The gradient of the slope for the c-fos regression is significantly greater than zero (ANOVA, F=14.82, p<0.01, n=12).

**Figure 6.**
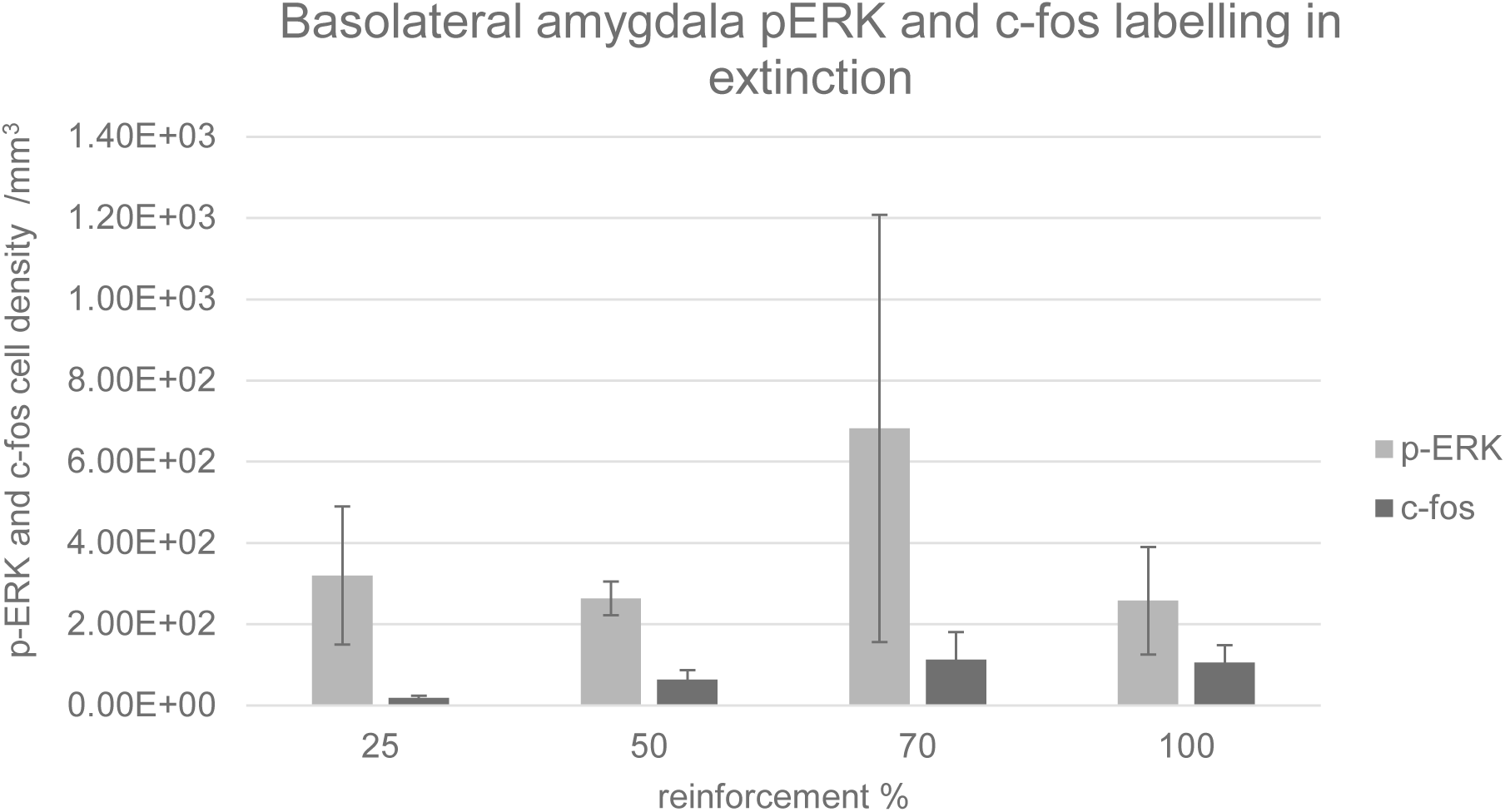
Density of p-ERK and c-fos labelled cells per mm^3^ in basolateral amygdala. Error bars indicate +/−1 standard error. There was no significant main effect of reinforcement group on p-ERK or c-fos cell density (ANOVA, p>0.05).

**Figure 7.**
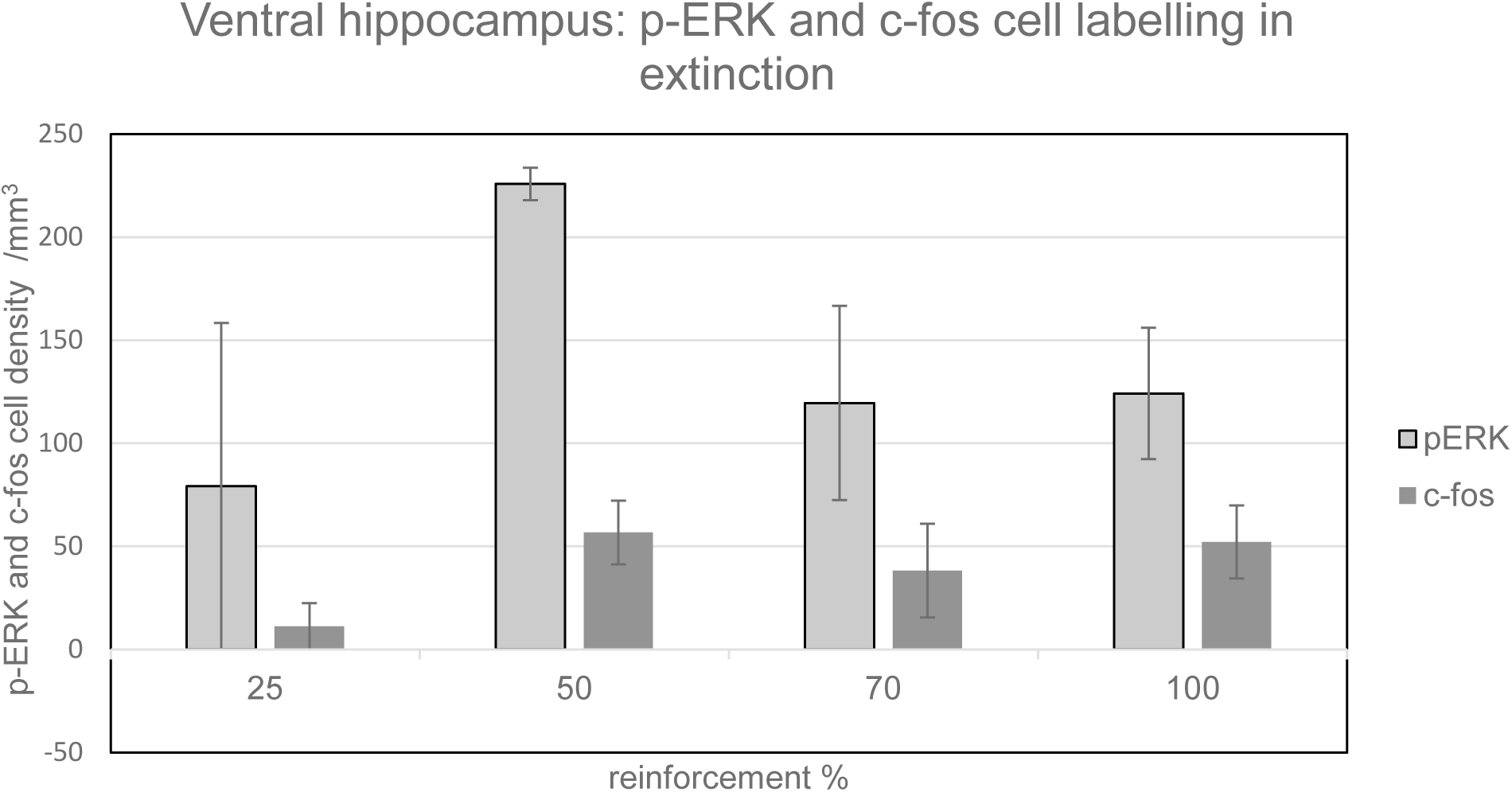
Density of p-ERK and c-fos labelled cells per mm^3^ in ventral hippocampus. Error bars indicate +/−1 standard error. There was no significant main effect of reinforcement group on p-ERK or c-fos cell density in a one-way ANOVA. Labelling in the 50% group was significantly greater than the 25%, 70% and 100% (t test, p<0.05, n=3 for each group).

**Figure 8.**
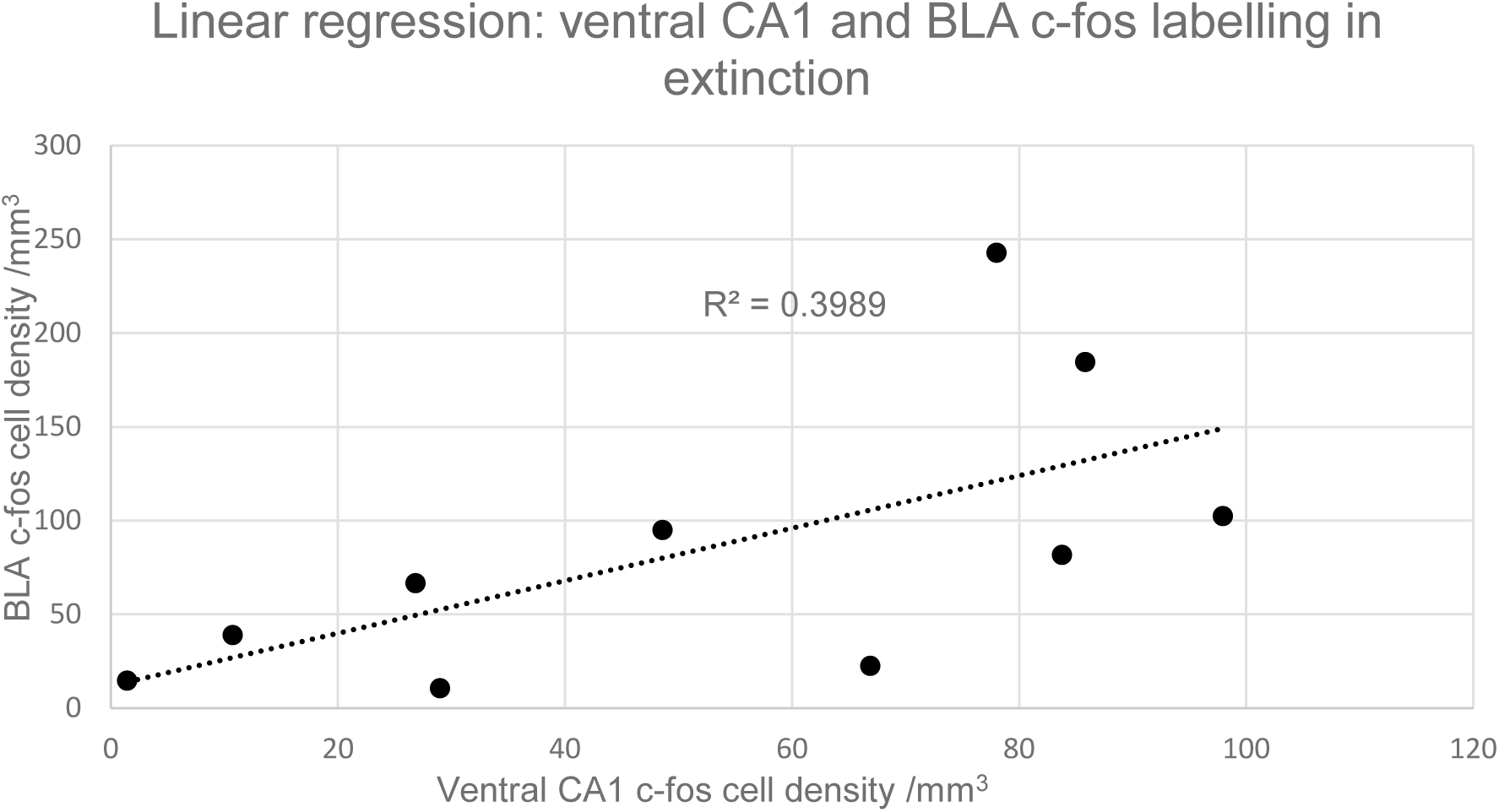
Graph of linear regression between mean c-fos cell density in ventral CA1 and BLA for all subjects. The gradient of the slope for the c-fos regression is significantly greater than zero (ANOVA, F=5.31, p<0.05); n=10.

## Results: Experiment 2

### Behaviour

Tone-evoked freezing in extinction was significantly greater in the 50% group than in the 100% (ANOVA, F=43.57, p<0.01, n=9, Figure 9). The PREE in Experiment 2 was noticeably more pronounced than in Experiment 1, despite the similar conditioning sequences (Figures 1 and 9). Other differences in experimental apparatus and procedure may account for the larger effect in Experiment 2: subjects in Experiment 2 were conditioned and extinguished in a larger, more open arena than the confined conditioning boxes in Experiment 1, and were in a post-surgical state, with electrodes in situ for both acquisition and extinction sessions.

**Figure 9.**
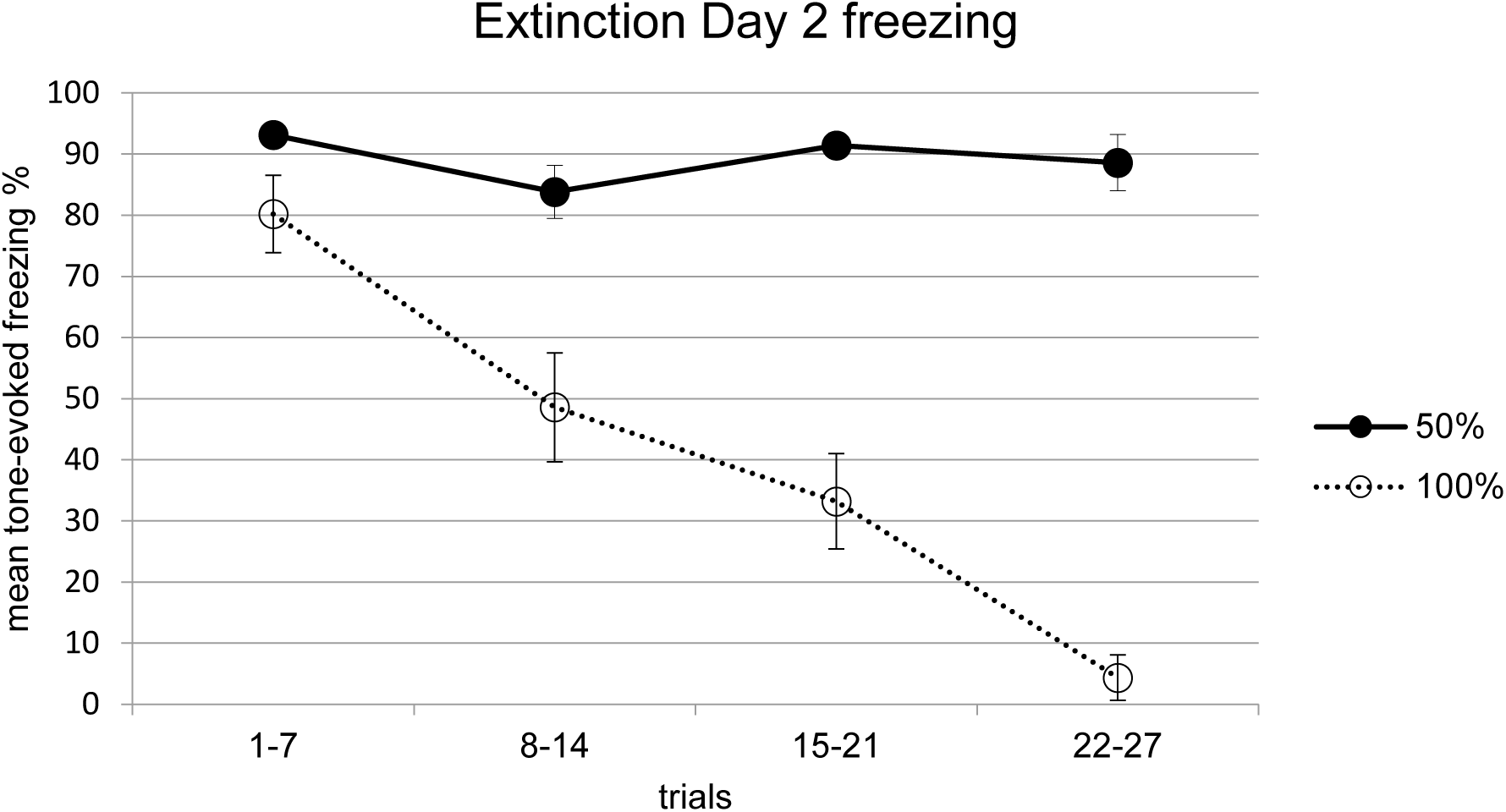
Freezing scores in extinction in Experiment 2. Freezing was significantly greater in the 50% than the 100% group (ANOVA, F=43.57, p<0.001, n=9). Error bars indicate +/−1 standard error.

### Electrophysiology

During acquisition (Day 1), CS tones evoked increased power in a high theta frequency band (8-10Hz, peak=9.3 Hz) of BLA LFPs in the 50% group (n=5, Figure 10). High theta (8-10Hz) responses were significantly greater during the CS tone compared to the pretone period in the 50% group (ANOVA, n=5, F=5.36, p<0.05, Figure 12). In the 100% group, the CS tone evoked evoked increases in LFP spectral power at lower theta frequencies (5-7Hz), although these increases were not significant (Figure 11). There were no significant tone-evoked increases in the 8-10Hz theta band in the 100% group (Figures 11 and 12). A CS tone-evoked spectral response in a high theta frequency band (8-10Hz) BLA was also observed in subiculum of 50% subjects during acquisition, although it did not reach statistical significance (ANCOVA, F=3.69, p=0.07, Figures 13 and 15). The peak of the 50% subiculum LFP spectral response (9.3Hz) was the same as that seen in BLA (Figures 10 and 13). There was an LFP spectral response in the subiculum to the CS tone in a lower theta frequency band (5-7Hz) in the 100% group, similar to that seen in BLA, but this response was not statistically significant (ANOVA, n=4, p>0.05; Figure 14).

**Figure 10.**
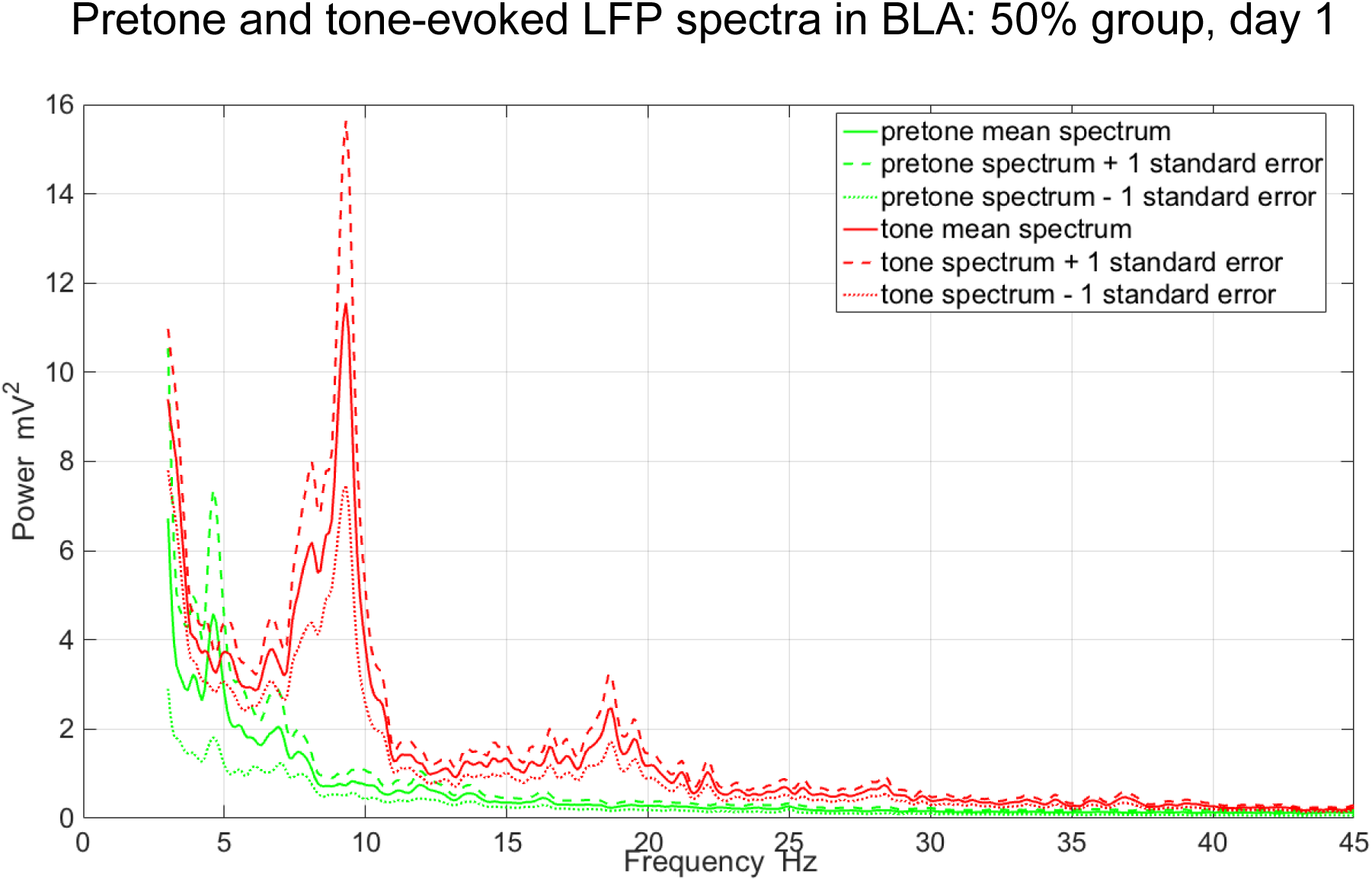
Pretone and tone-evoked spectral power of local field potentials (LFPs) in BLA in the 50% group during acquisition. The CS tone evoked mean increases in LFP spectral power in the theta frequency range (peak=9.3 Hz, n=5).

**Figure 11.**
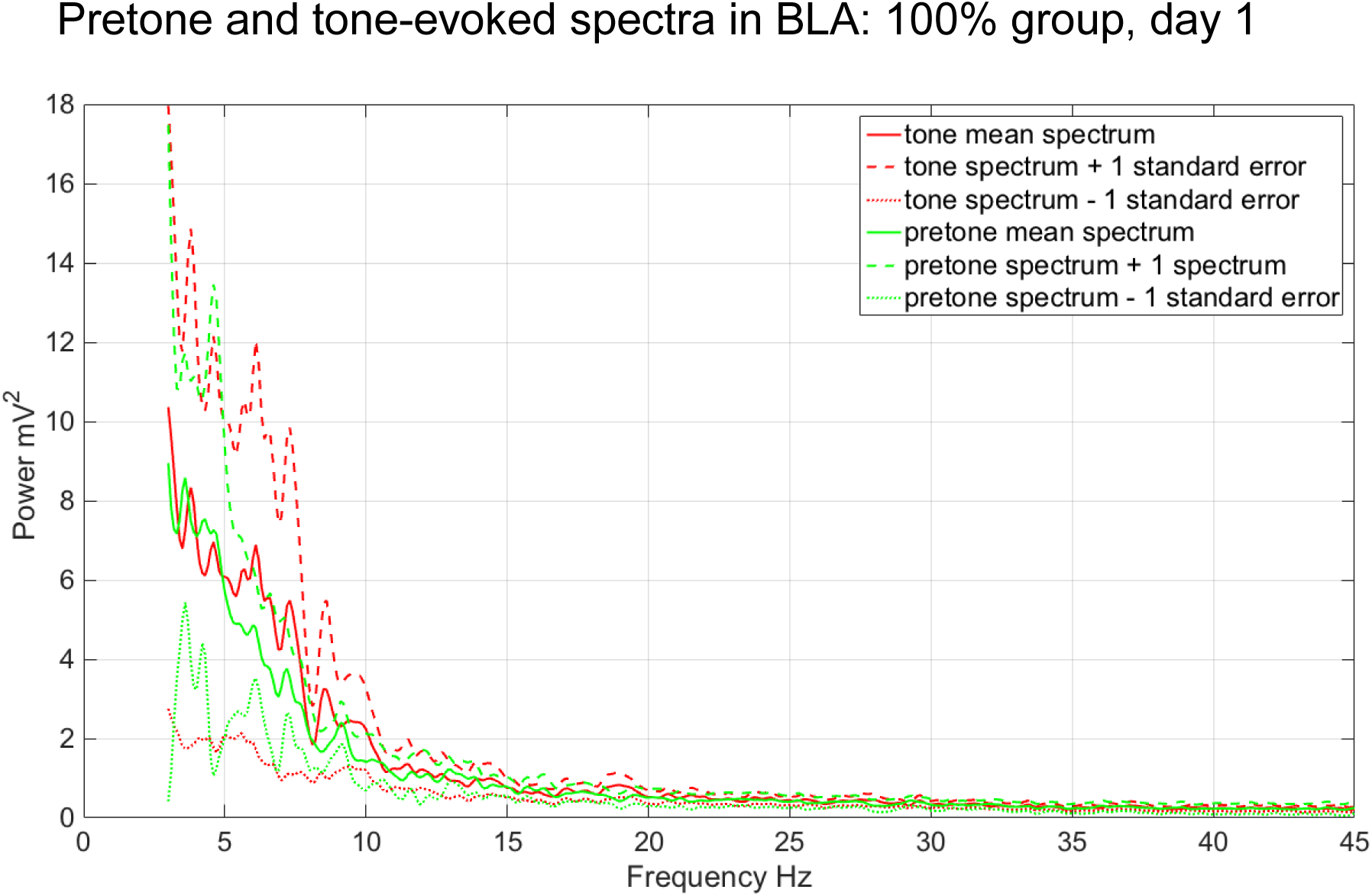
Pretone and tone-evoked LFP spectral power in BLA in the 100% group during acquisition (n=4).

**Figure 12.**
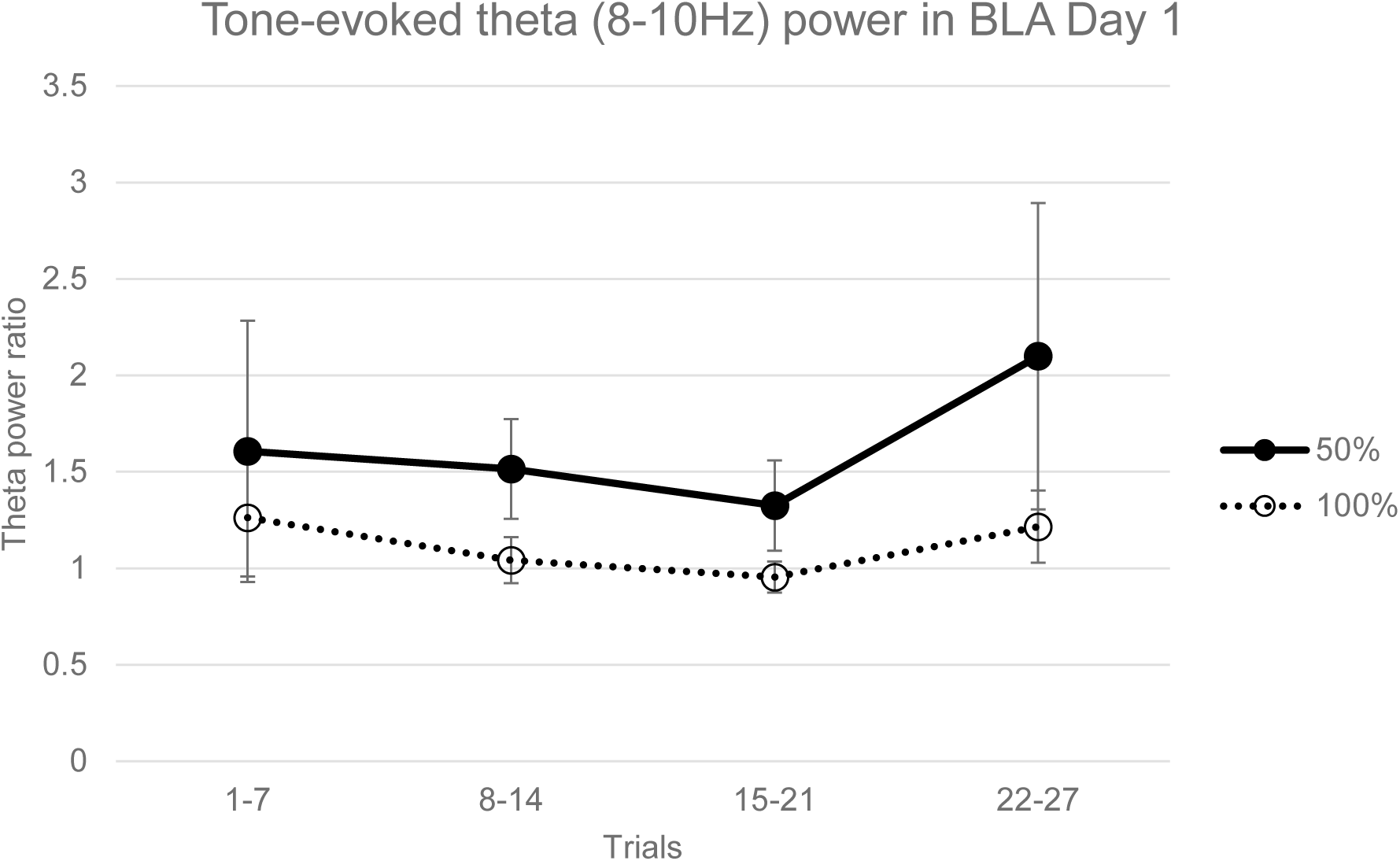
Tone-evoked normalized LFP theta (8-10 Hz) power in BLA during acquisition. Theta power during the CS tone was significantly greater than the pretone period in the 50% group (ANOVA, F=5.36, p<0.05), but not in the 100% group (ANOVA, n=4, p>0.05). Error bars indicate +/−1 standard error.

**Figure 13.**
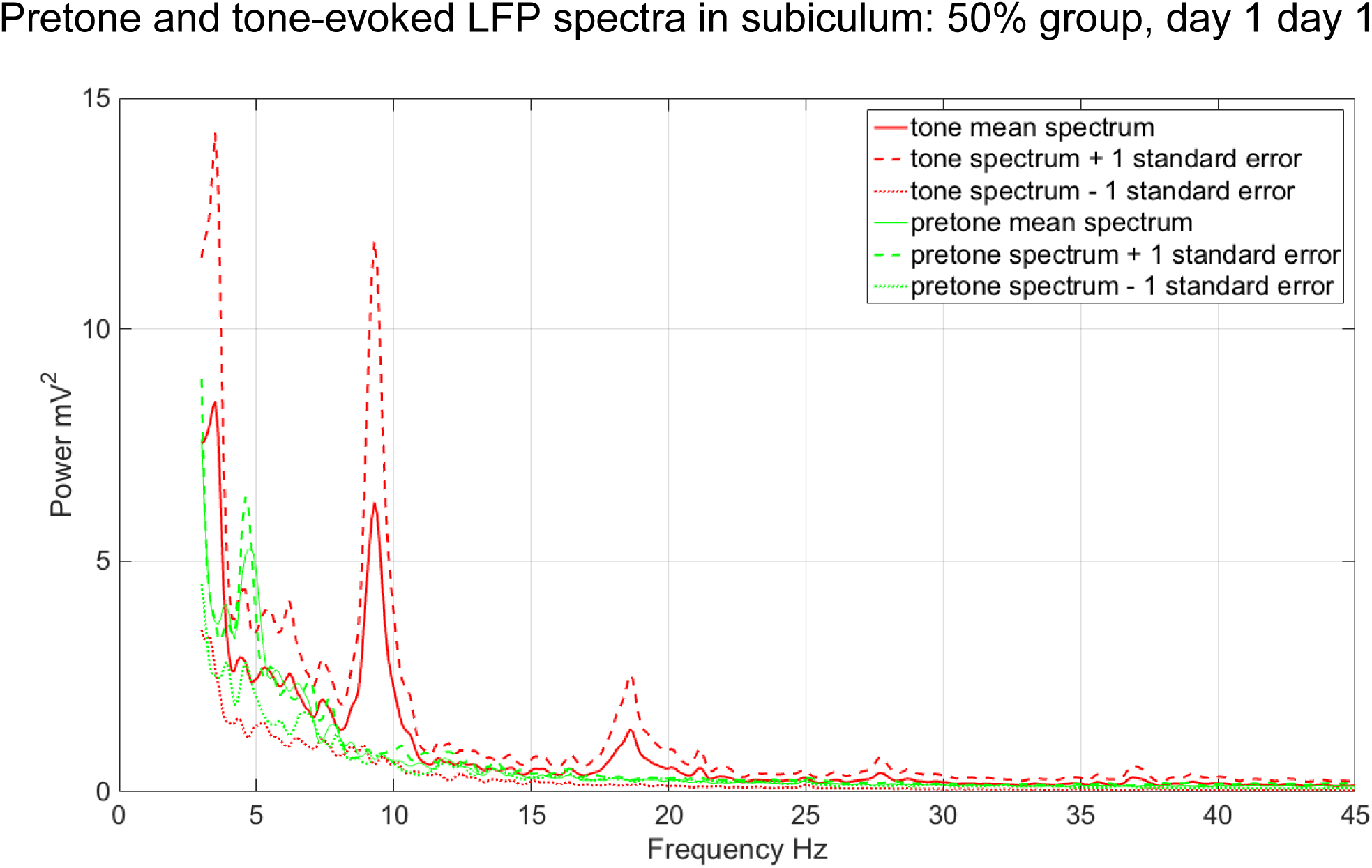
Pretone and tone-evoked LFP spectral power in subiculum in the 50% group during acquisition (n=3). The CS tone evoked mean increases in LFP spectral power in the theta (peak=9.3 Hz) and delta (peak=3.5Hz) LFP frequencies (n=3).

**Figure 14.**
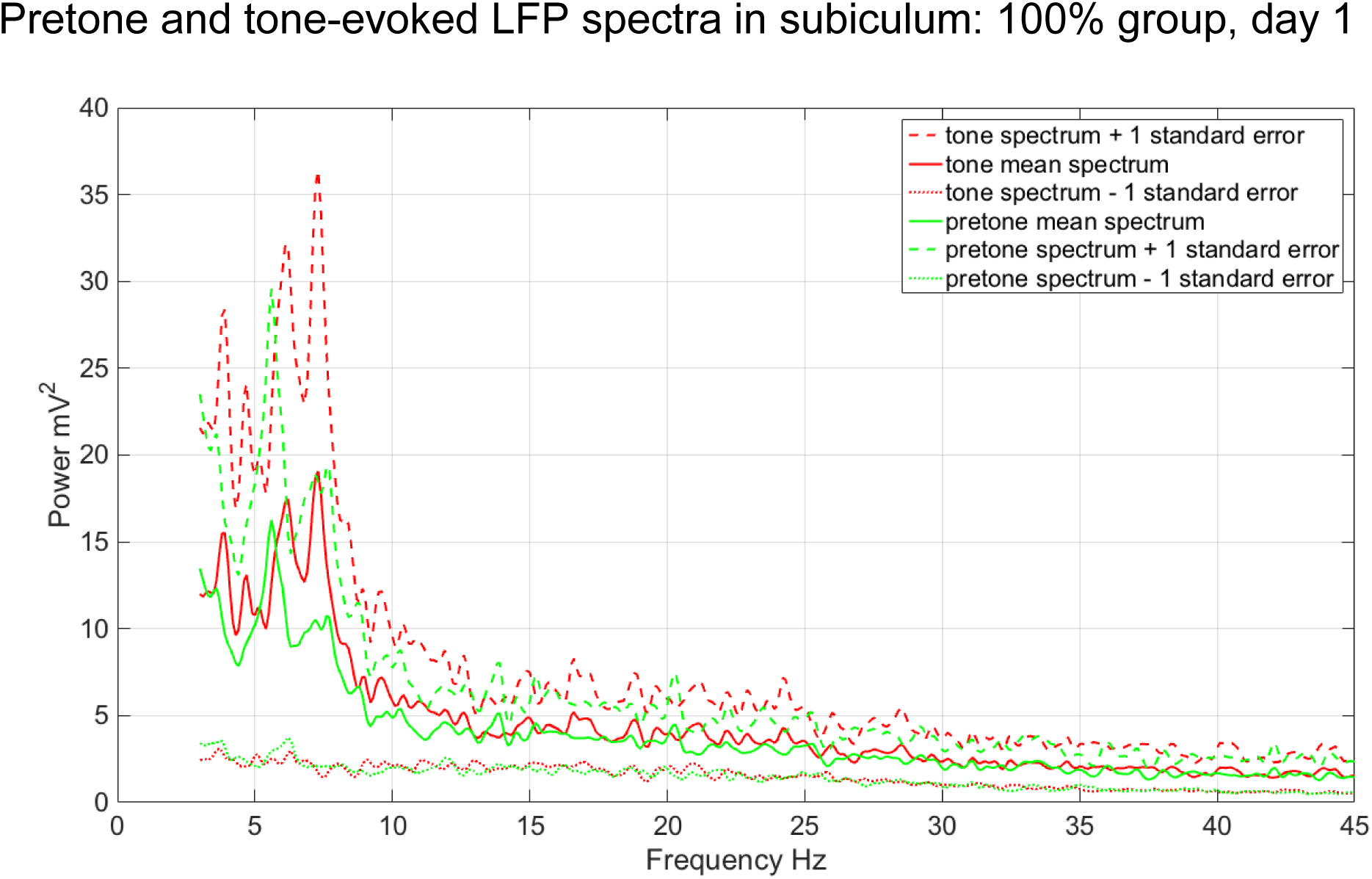
Pretone and tone-evoked LFP spectral power in subiculum in the 100% group during acquisition. The CS tone evoked mean increases in LFP spectral power in the theta frequency range (peak=7.3 Hz, n=2).

**Figure 15.**
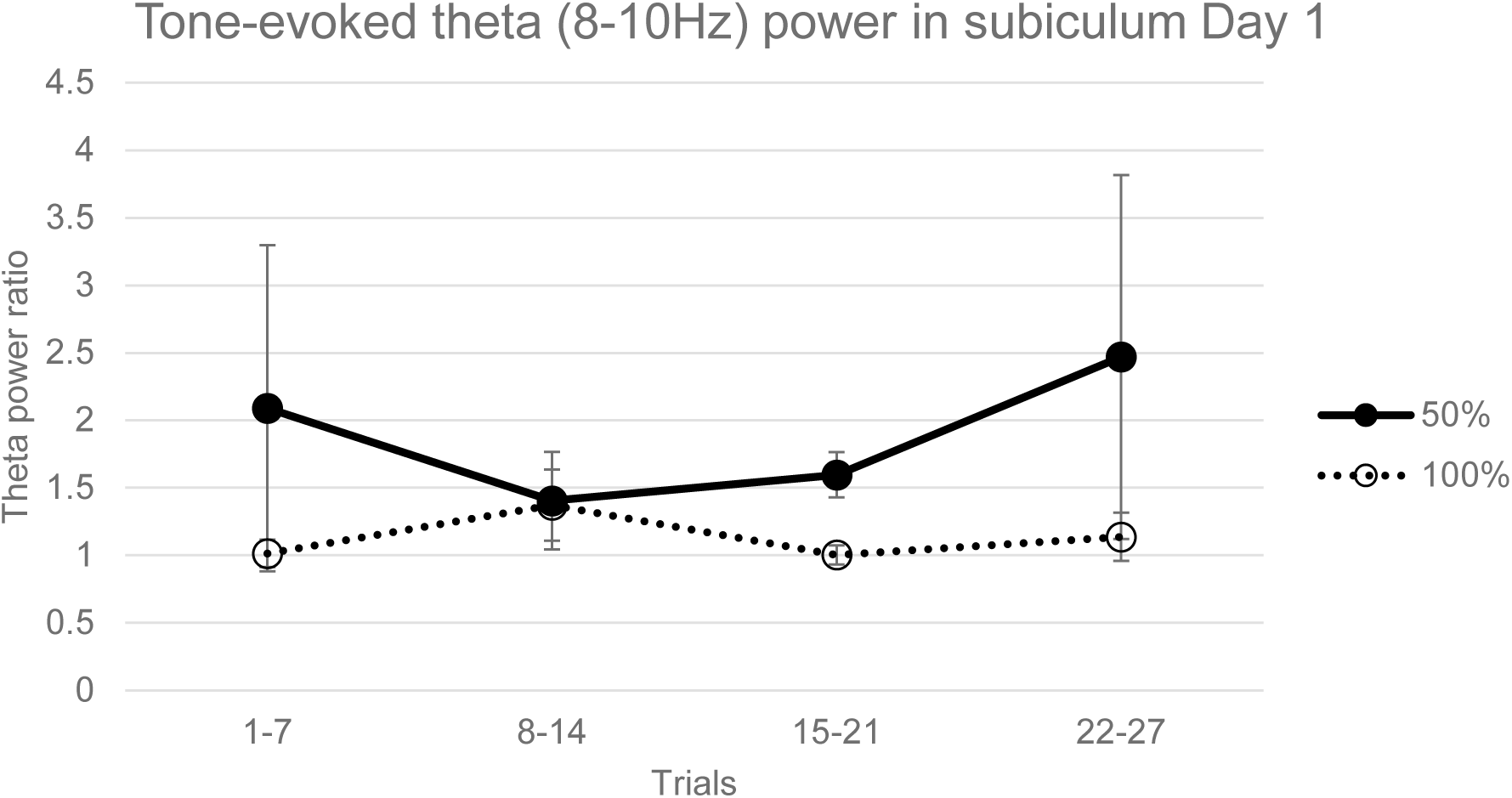
Tone-evoked normalized theta (8-10 Hz) power in subiculum in acquisition. Theta power during the CS tone was not significantly greater than the pretone period in the 50% group (ANOVA, n=5, F=3.69, p=0.07), or in the 100% group (ANOVA, n=4, p>0.05). Error bars indicate +/−1 standard error.

During extinction (Day 2), a significant theta (8-10Hz) spectral response to the CS tone was seen in the 50% subjects in BLA (ANCOVA, F=11.01, p<0.01, Figures 16 and 18), with a similar peak (9.4Hz) to that seen on Day 1 (Figure 10). In 100% subjects, non-significant increases in LFP spectral power in BLA were seen in 8-10Hz and 12-14Hz frequency bands (ANOVA, n=4, p>0.05; Figures 17 and 18). A significant CS tone-evoked theta (8-10Hz) LFP response, with a similar peak (9.4Hz), was also seen in the subiculum during extinction (ANCOVA, F=6.51, p<0.05, Figures 19 and 21). In the 100% group, a non-significant LFP spectral response in the subiculum to the CS tone was seen in a 12-14Hz frequency band (ANOVA, n=4, p>0.05; Figures 20 and 21) similar to that seen in BLA (Figure 17).

**Figure 16.**
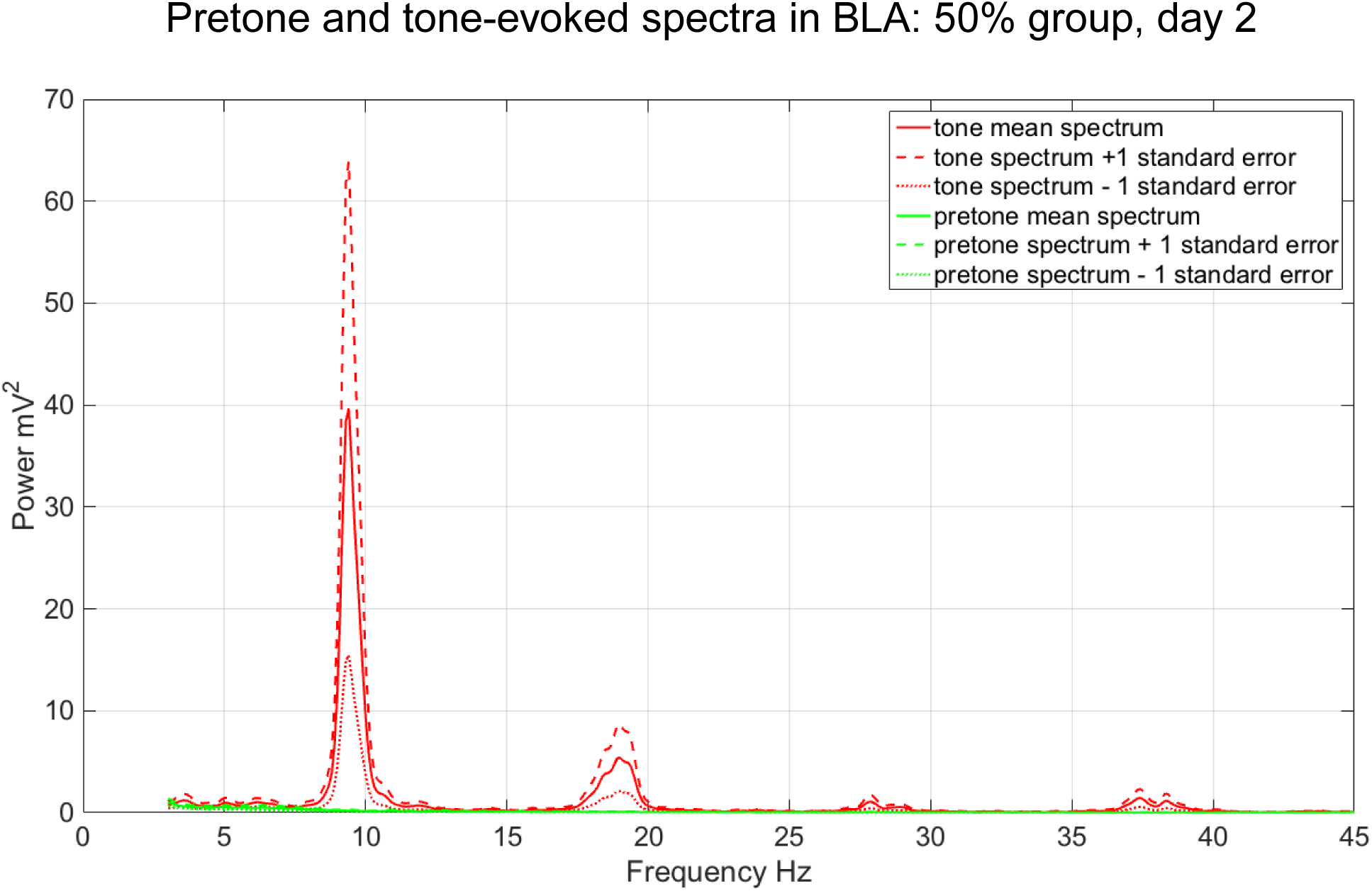
Pretone and tone-evoked LFP spectral power in BLA in the 50% group during extinction (n=5). The CS tone evoked mean increases in LFP spectral power in theta LFP frequencies (peak=9.4 Hz, n=5).

**Figure 17.**
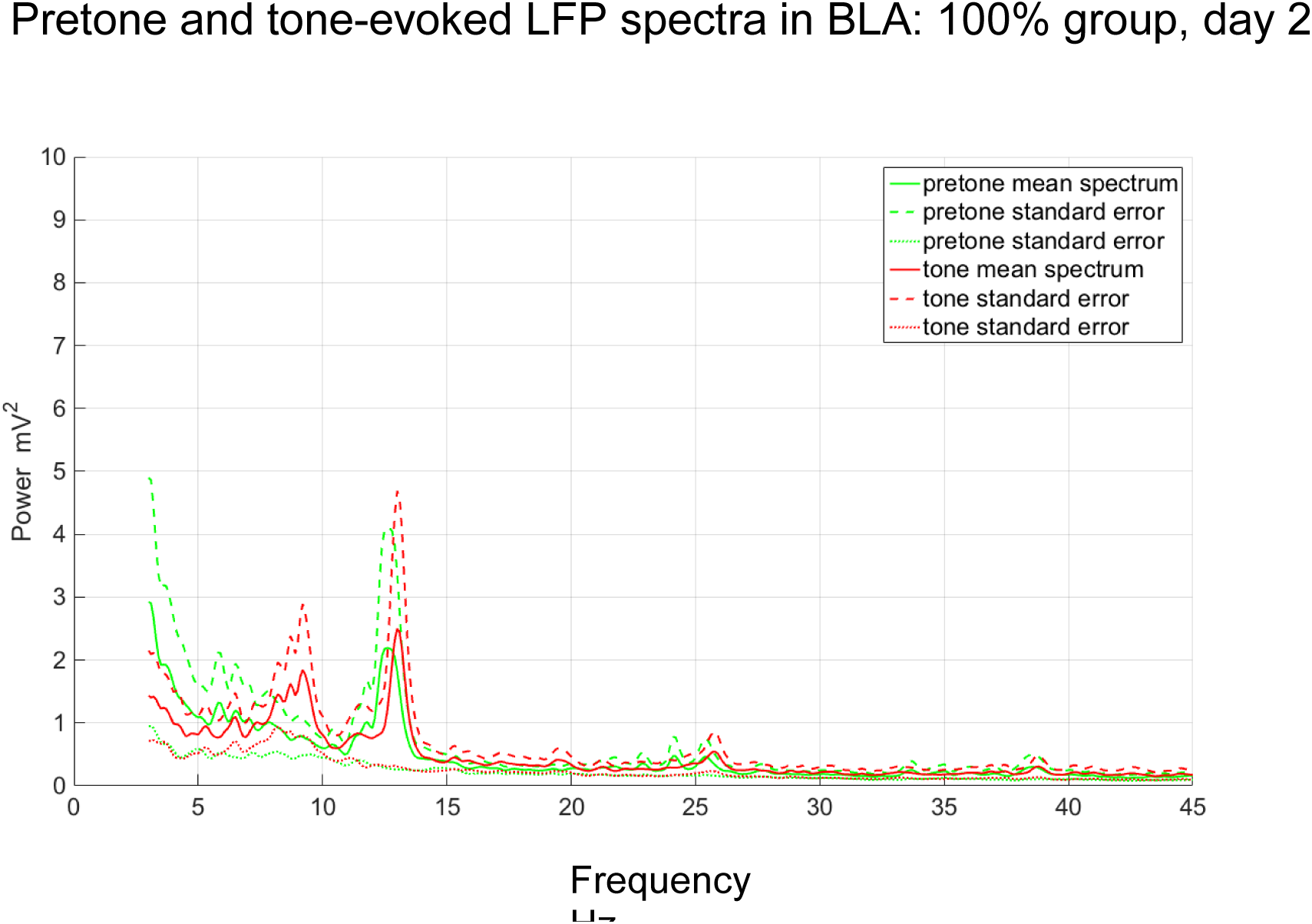
Pretone and tone-evoked LFP spectral power in BLA in the 100% group during extinction. The CS tone evoked mean increases in LFP spectral power in the theta frequency range (peaks=9.2 Hz, 13.0Hz, n=5).

**Figure 18.**
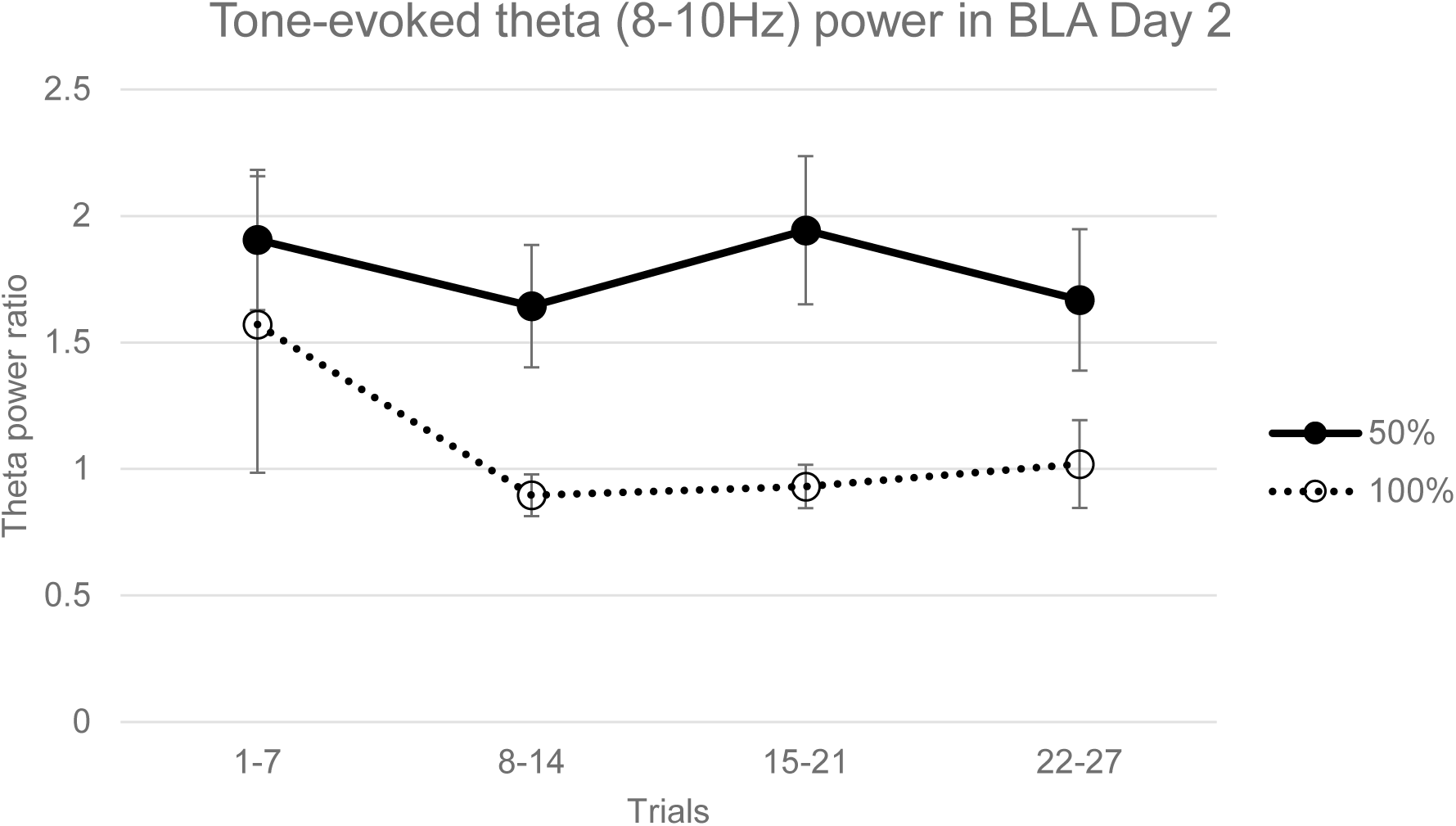
Tone-evoked normalized theta (8-10 Hz) power in BLA in extinction. Theta power during the CS tone was significantly greater in the 50% group compared to the 100% group (ANOVA, F=11.01, p<0.01.

**Figure 19.**
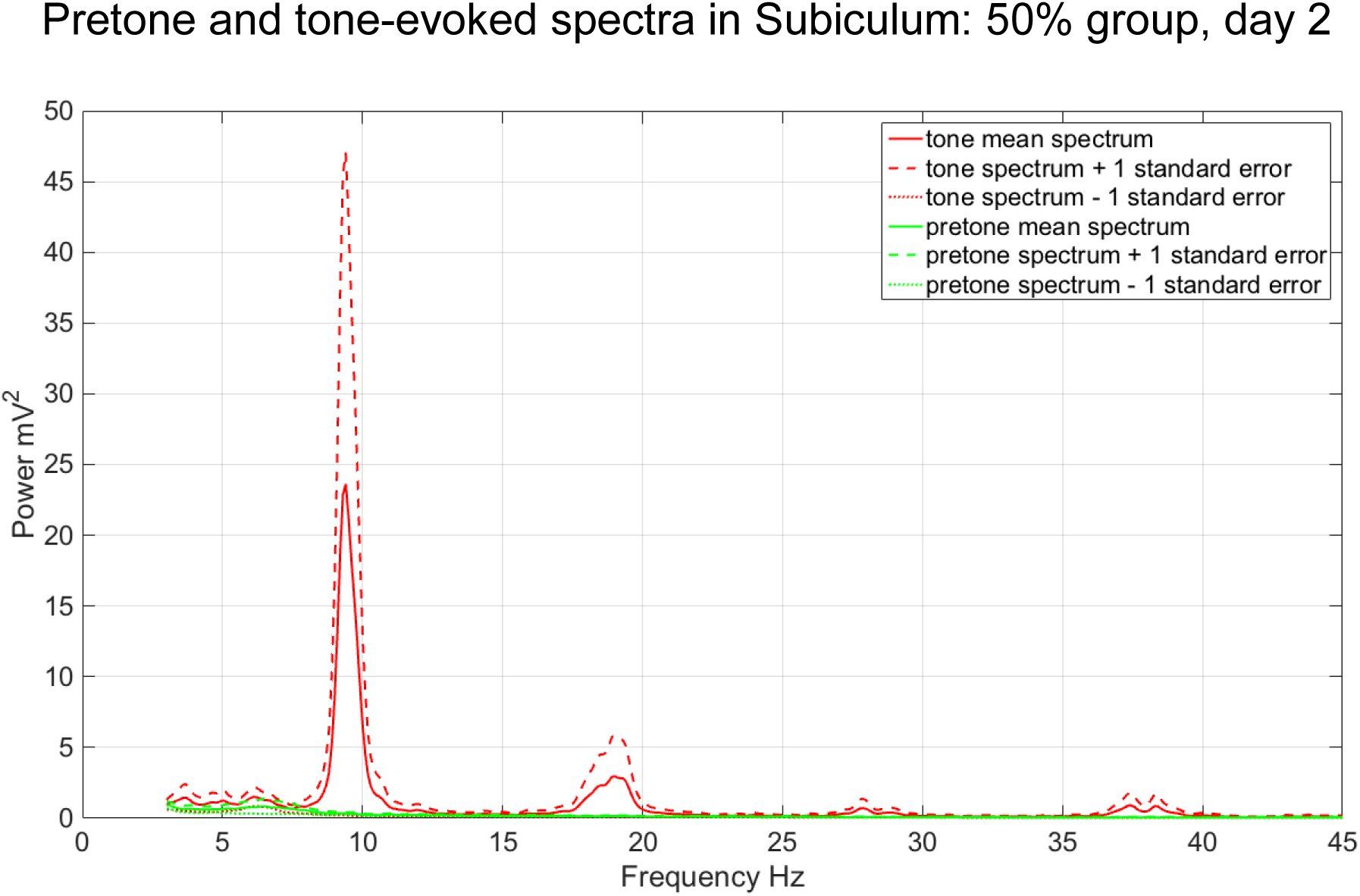
Pretone and tone-evoked LFP spectral power in the subiculum in the 50% group during extinction (n=3). The CS tone evoked mean increases in LFP spectral power in theta LFP frequencies (peak=9.4 Hz, n=5).

**Figure 20.**
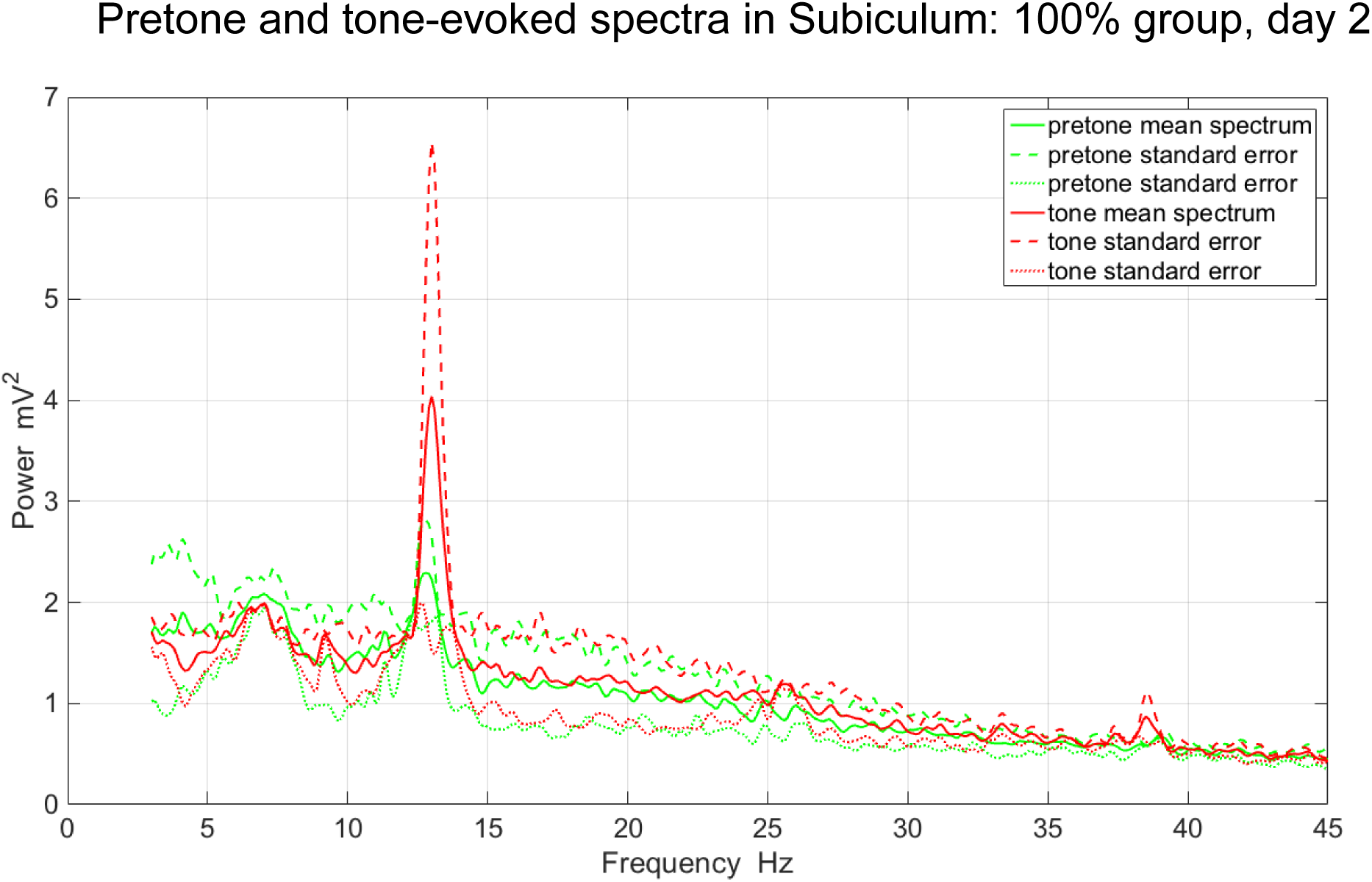
Pretone and tone-evoked LFP spectral power in subiculum in the 100% group during acquisition (n=2). The CS tone evoked mean increases in LFP spectral power in the theta frequency range (peak=13.0Hz, n=2).

**Figure 21.**
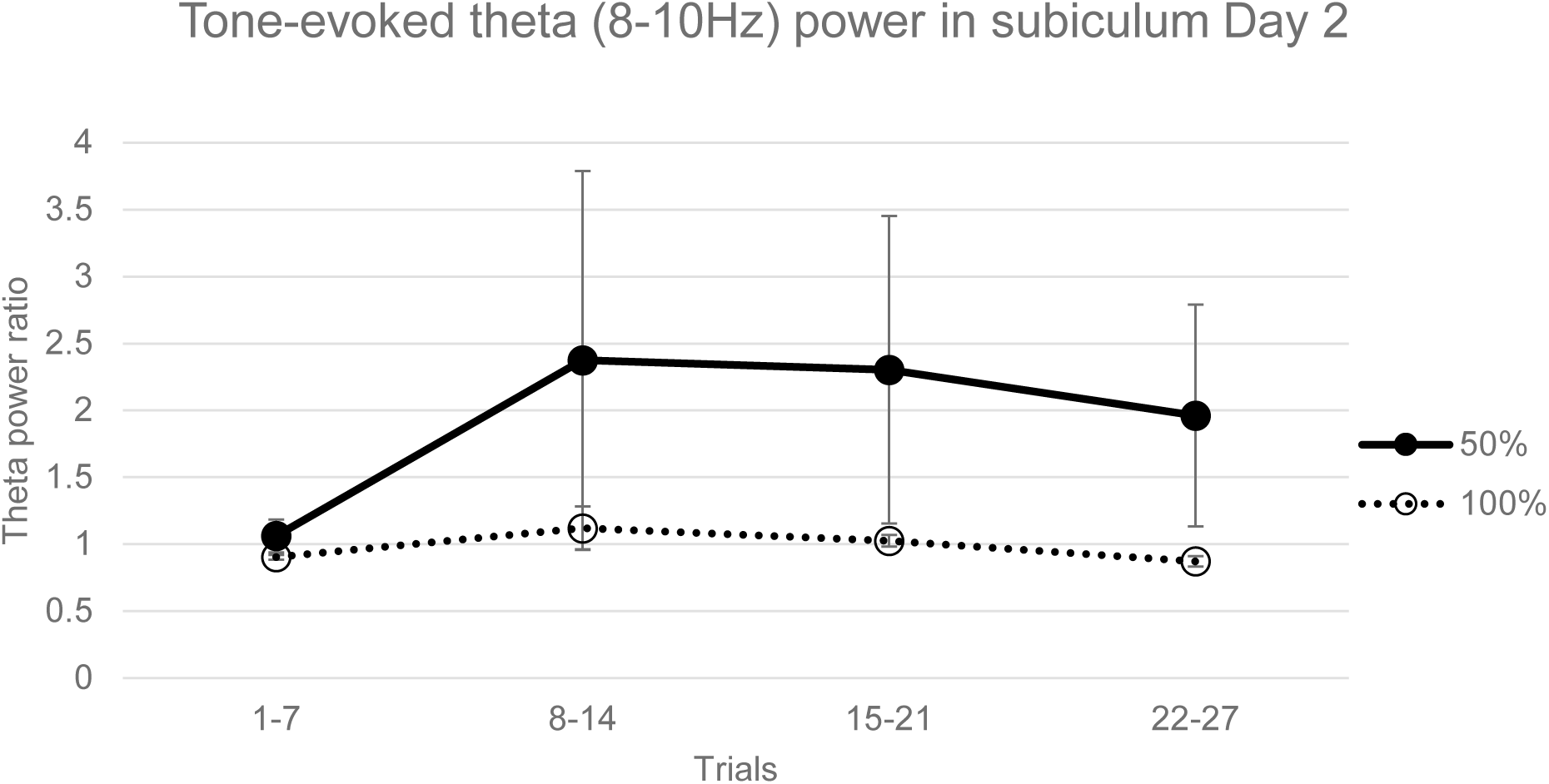
Tone-evoked normalized theta (8-10 Hz) power of local field potentials (LFPs) in subiculum in acquisition. Theta power during the CS tone was significantly greater than the pretone period in the 50% group (ANOVA, F=6.51, p<0.05), but not in the 100% group. Error bars indicate +/−1 standard error.

Linear coherence between BLA and subiculum LFP channels during the acquisition session was greater in the 50% than the 100% group across a broad frequency range and in both pretone and tone periods (Figure 22), with a significant difference in the theta (8-10Hz) frequency band (ANCOVA, f=10.17, p<0.01, Figure 23). In addition, mean BLA-subiculum coherence was greater during CS tone periods compared to pretone periods across the whole frequency range in the 50% group, but not in the 100% group (Figure 22) with a significant difference in the 8-10Hz frequency range (ANCOVA, n=9, F=8.91, p<0.01, Figure 23). In the extinction session, a similar pattern of coherence was observed (Figure 24), with the 50% subjects having significantly greater BLA-subiculum coherence than 100% subjects in a 8-10Hz frequency band (ANCOVA, F=17.01, p<0.01, Figure 25) and with tone coherence being significantly greater than pretone in the 50% group, but not in the 100% group (ANCOVA, F=27.03, p<0.001, Figure 25).

**Figure 22.**
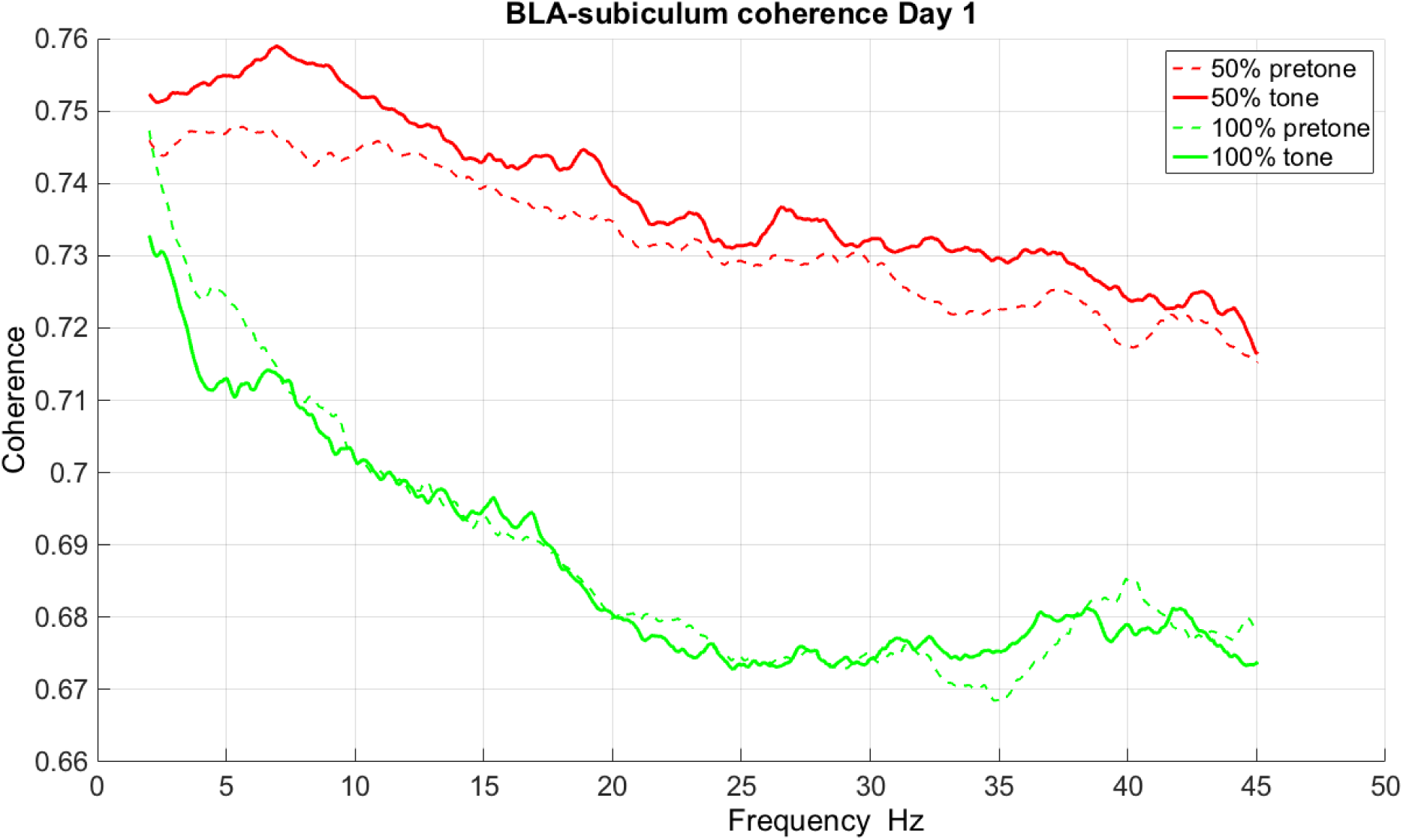
Coherence between local field potentials in BLA and subiculum during acquisition. Coherence is shown for pretone and tone periods in both 50% and 100% groups.

**Figure 23.**
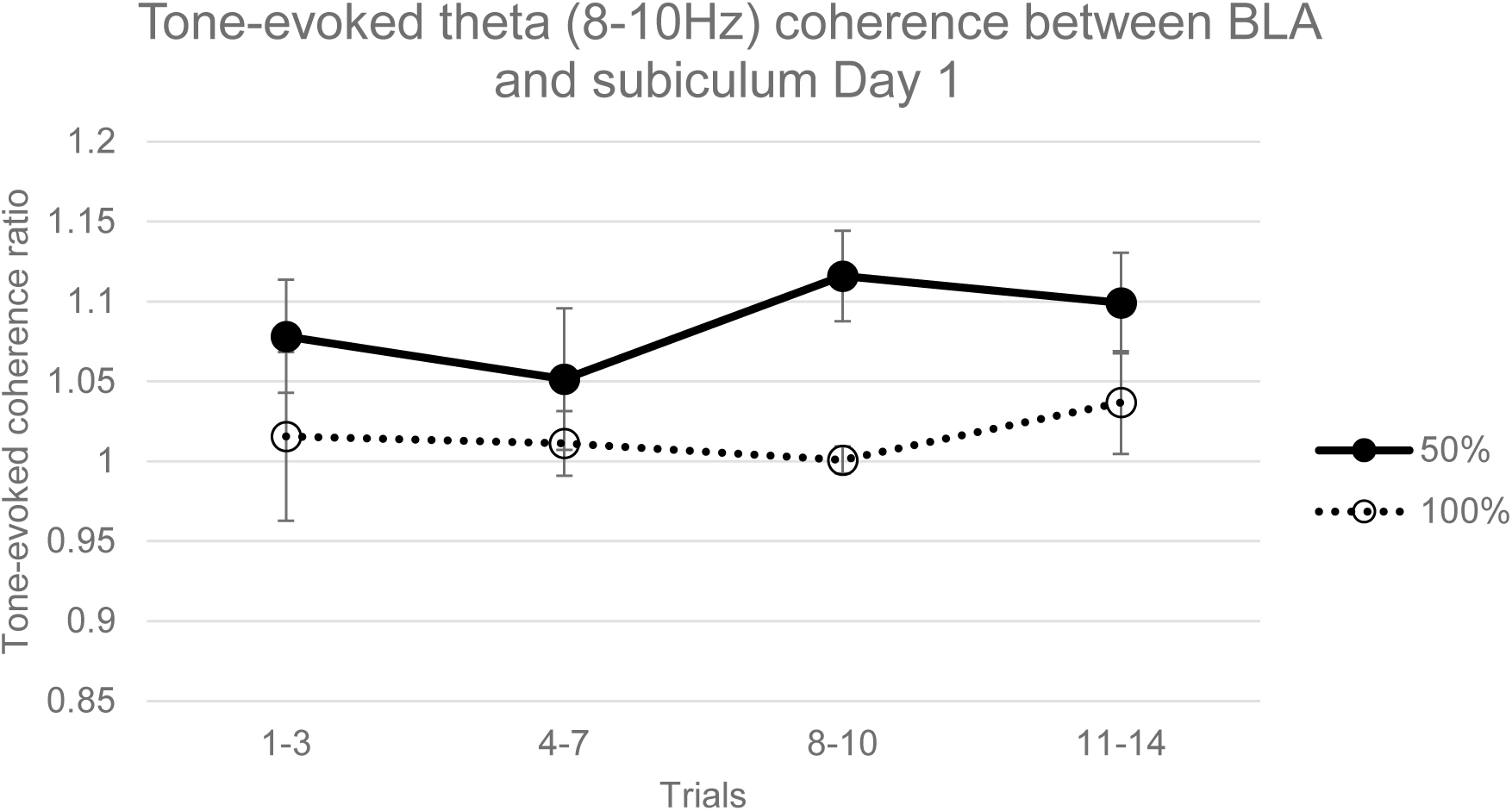
Tone-evoked theta (8-10Hz) coherence between BLA and subiculum LFPs during acquisition in 50% and 100% groups. Coherence in the 50% tone period was significantly greater than both the 50% pretone period (ANOVA, F=8.91, p=0.01) and the 100% tone period (ANOVA, F=10.17, p<0.01). Error bars indicate +/−1 standard error.

**Figure 24.**
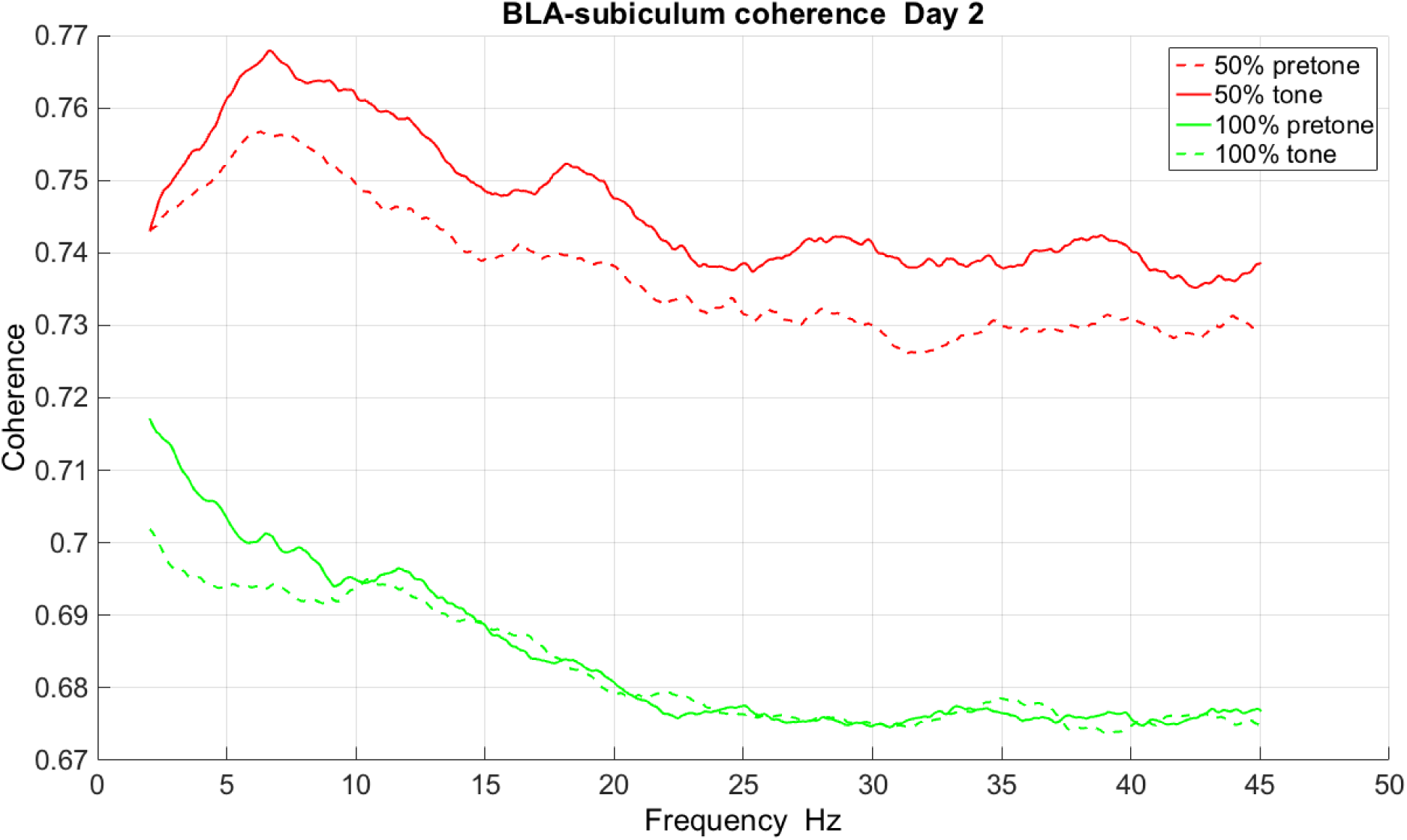
Coherence between local field potentials in BLA and subiculum during extinction. Coherence is shown for pretone and tone periods in both 50% and 100% groups.

**Figure 25.**
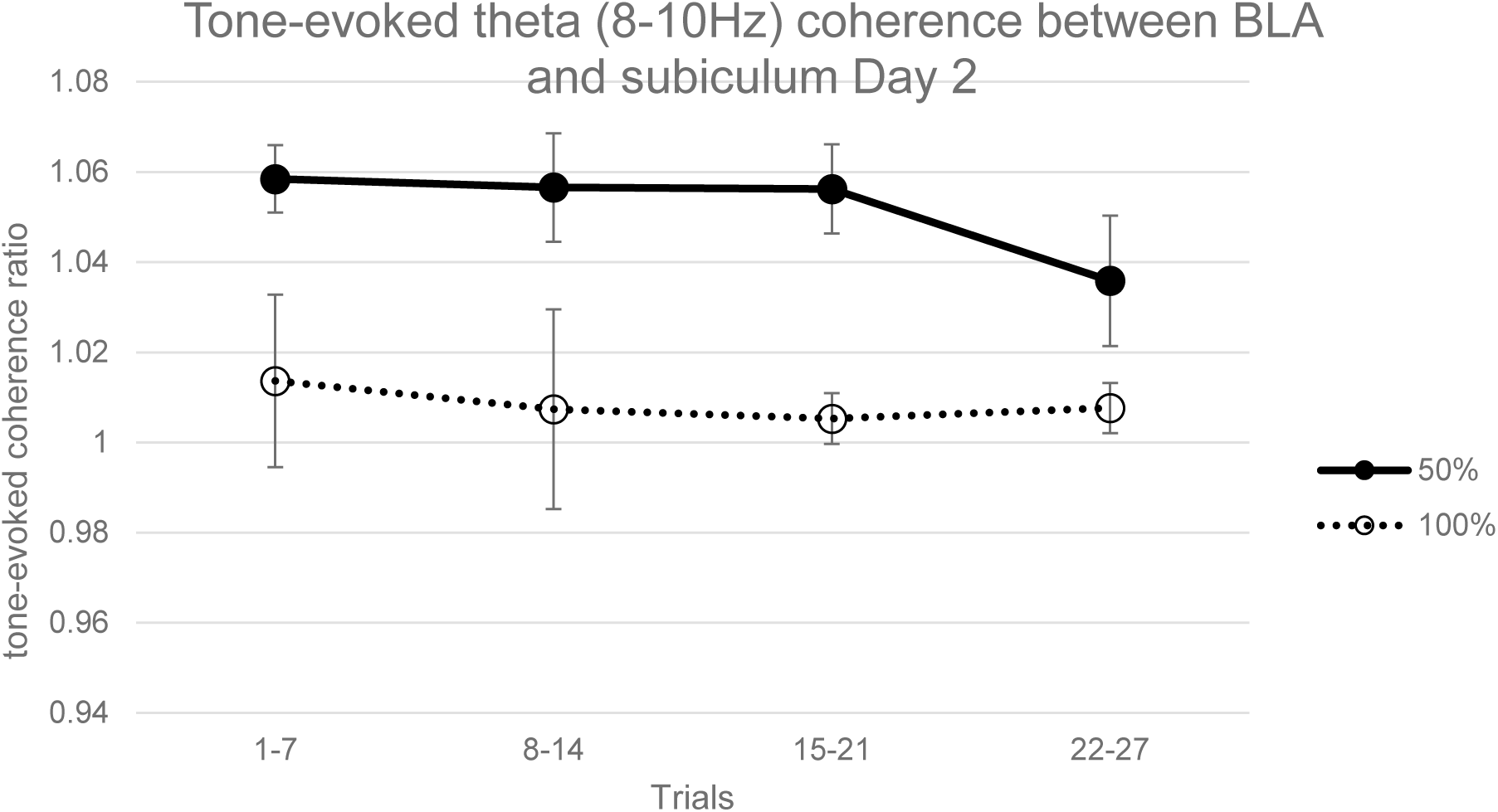
Tone-evoked theta (8-10Hz) coherence between BLA and subiculum LFPs during extinction in 50% and 100% groups. Coherence in the 50% tone period was significantly greater than the 50% pretone period (ANOVA, F=27.03, p<0.001) and the 100% tone period (ANOVA, F=17.01, p<0.01). Error bars indicate +/−1 standard error.

A Granger Causality analysis of acquisition LFPs revealed no significant differences between 50% and 100% subjects or between pretone and tone periods in either the BLA=>subiculum direction (Figure 26) or subiculum=>BLA direction (Figure 27). This is in contrast to the coherence analysis of the same data, which showed significant differences between 50% and 100% groups and between tone and pretone periods during acquisition (Figures 22 and 23). In the extinction session, Granger Causality in the BLA=>subiculum direction was significantly greater in the 50% group compared to the 100% group in both pretone (ANOVA, F=4.74, p<0.05) and tone (ANOVA, F=4.96, p<0.05) periods (Figure 28). In the subiculum=>BLA direction, there was a trend towards greater Granger Causality in the 50% group in the pretone (ANOVA, F=4.1, p=0.066) and tone (ANOVA, F=4.02, p=0.068) periods (Figure 29). In both BLA=>subiculum and subiculum=>BLA directions there were no significant differences between pretone and tone periods in extinction, in contrast to the significant coherence differences between pretone and tone periods seen in the 50% group (Figures 24 and 25).

**Figure 26.**
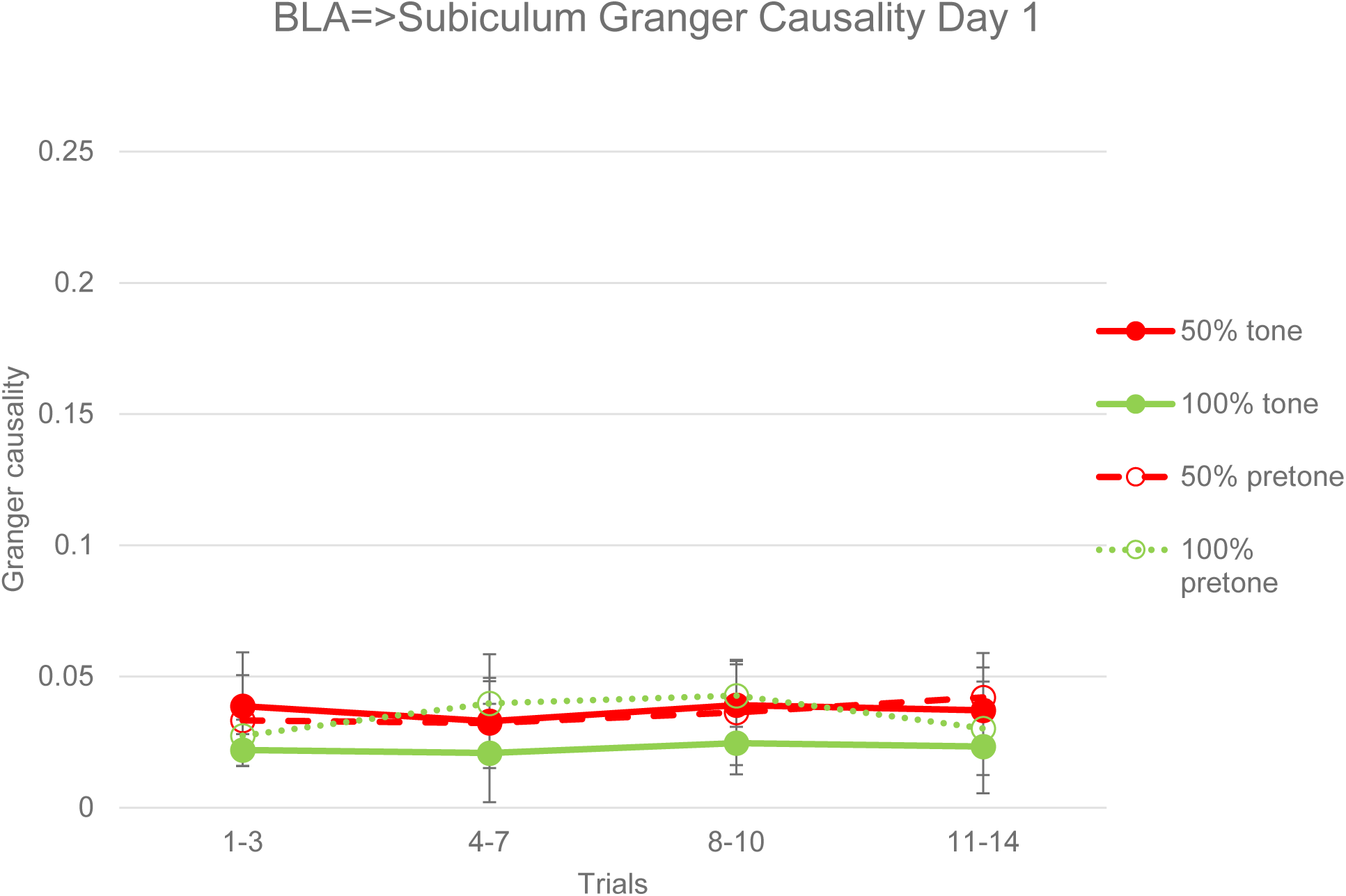
Granger Causality from BLA to subiculum LFPs in pretone and tone periods during acquisition. There were no significant differences in Granger Causality between 50% and 100% groups or between pretone and tone periods in either group. Error bars indicate +/−1 standard error.

**Figure 27.**
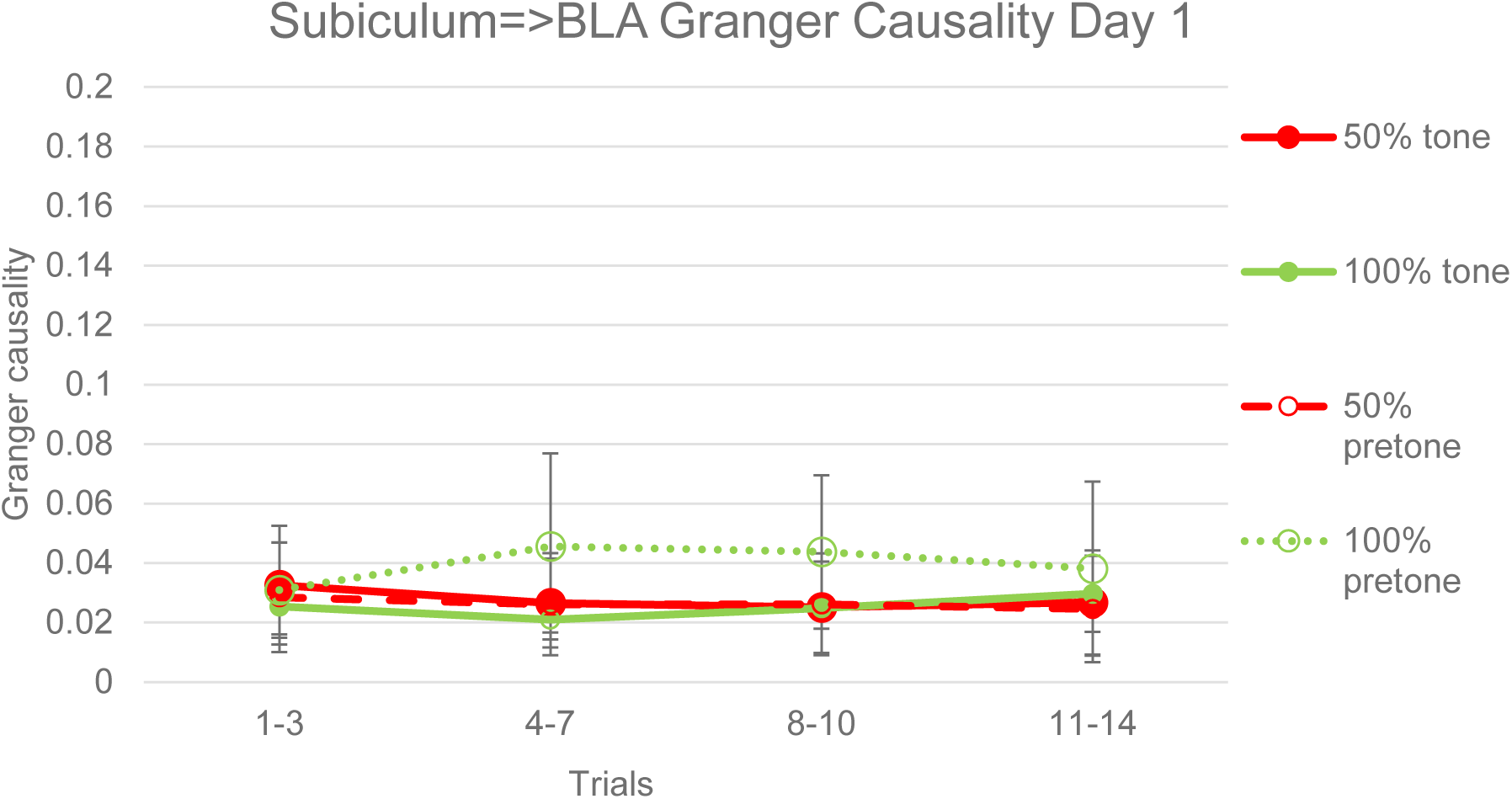
Granger Causality from subiculum to BLA LFPs in pretone and tone periods during acquisition. There were no significant differences in Granger Causality between 50% and 100% groups or between pretone and tone periods in either group. Error bars indicate +/−1 standard error.

**Figure 28.**
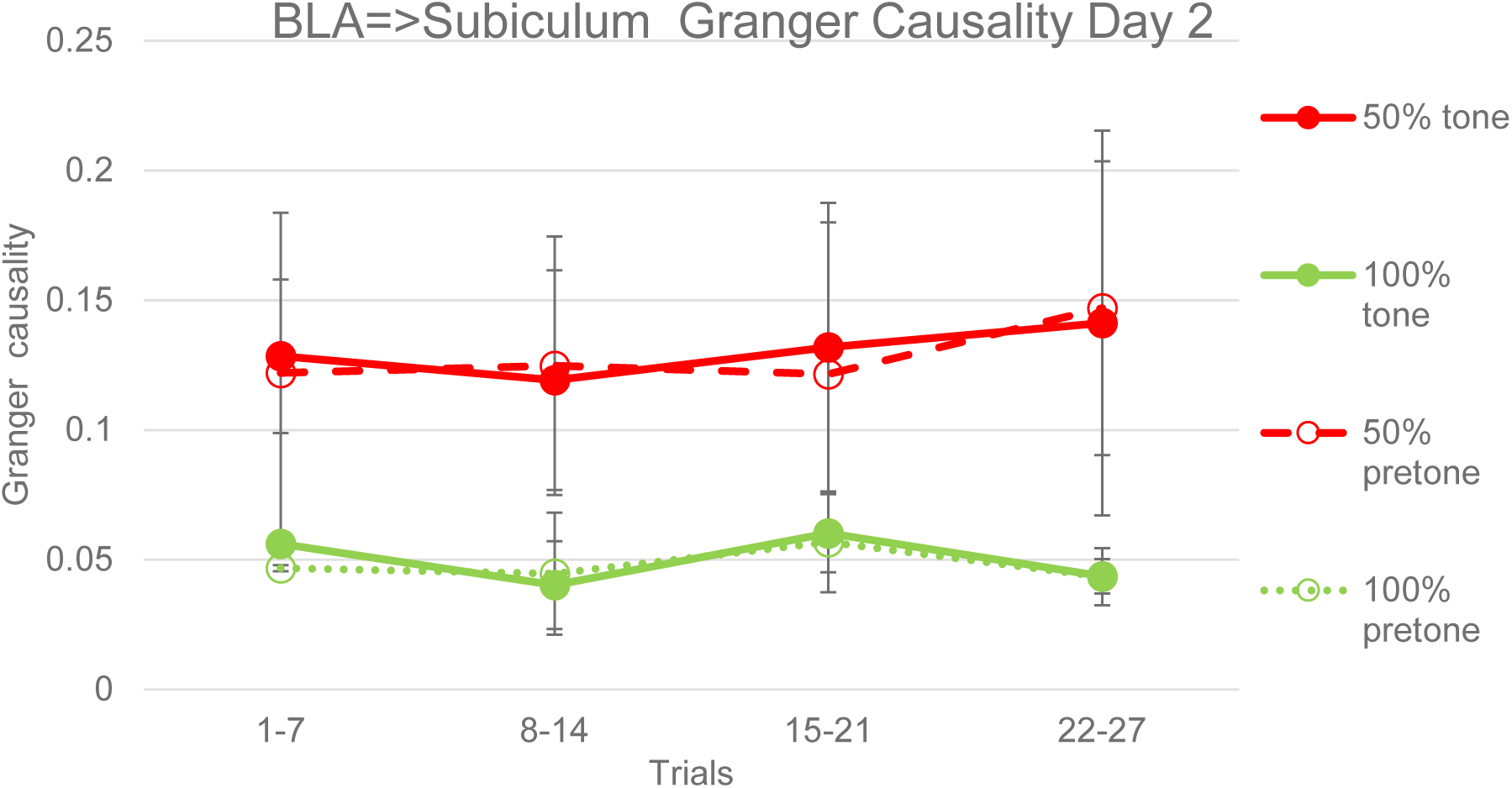
Granger Causality from subiculum to BLA LFPs in pretone and tone periods during acquisition. There were significant differences between the 50% and 100% groups in both pretone (ANOVA, F=4.74, p<0.05) and tone (ANOVA, F=4.96, p<0.05) periods. Error bars indicate +/−1 standard error.

**Figure 29.**
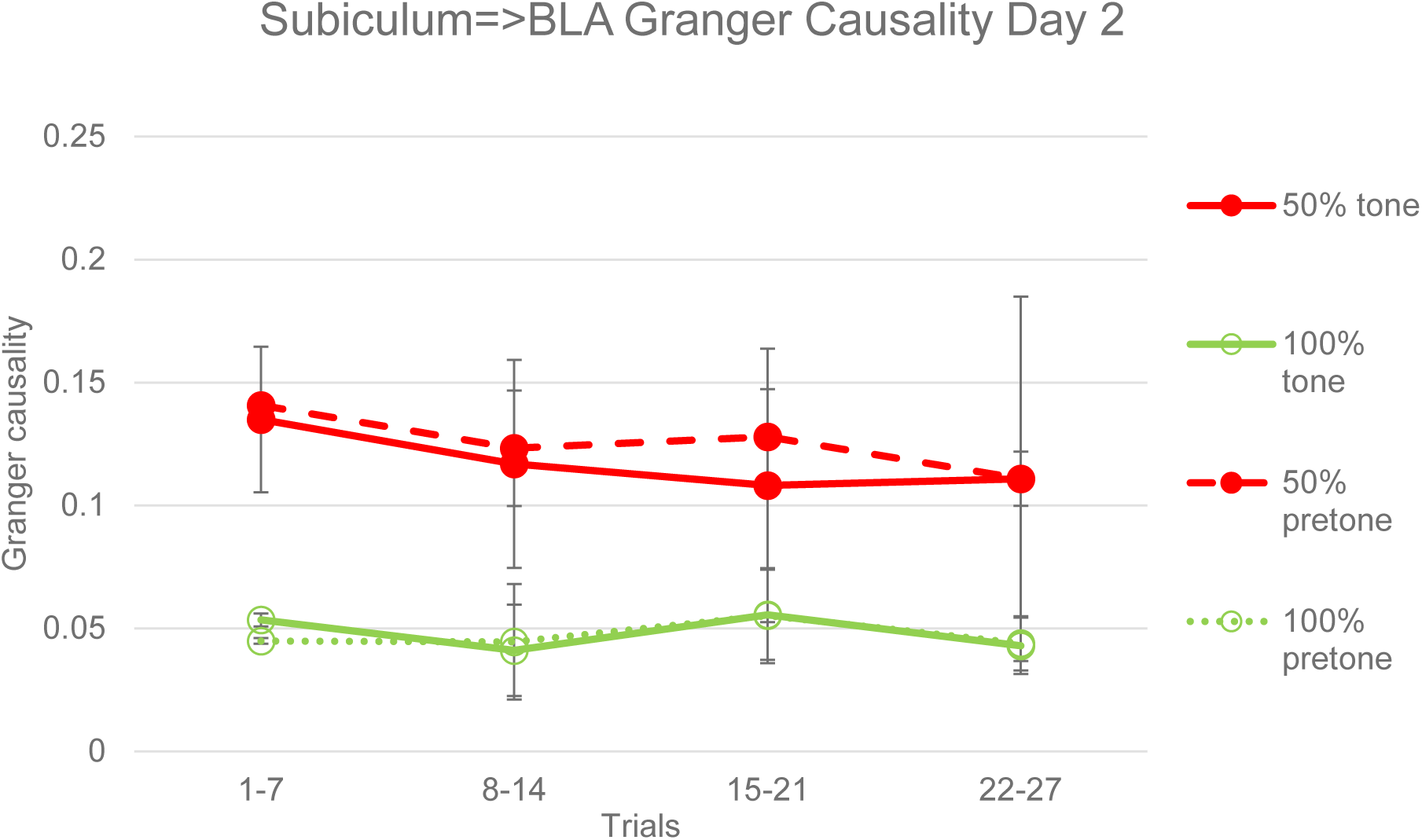
Granger Causality from subiculum to BLA LFPs in pretone and tone periods during acquisition. There were no overall significant differences between the 50% and 100% groups in either pretone (ANOVA, F=4.1, p=0.066) or tone (ANOVA, F=4.02, p=0.068) periods. Error bars indicate +/−1 standard error.

## Discussion

The significant behavioural PREE observed in the 50% reinforcement fear conditioning group (Figures 1, 2 and 9) is difficult to explain with standard associative learning theories (Rescorla and Wagner, 1972) or with rate estimation theory (Gallistel and Gibbon, 2000), given that the 50% and 100% groups had an identical number and rate of US presentations. However, the entropy-like, inverted U distribution of conditioned responding against reinforcement probability, which was also reported by a previous study of the PREE using a range of probabilities (Grant and Schipper 1952), is consistent with a Bayesian interpretation, which proposes that the phenomenon is driven by stimulus entropy or uncertainty (Courville et al. 2006). A Bayesian inferential account is also consistent with the observation that random (unpredictable) 50% reinforcement schedules reliably elicit greater resistance to extinction than regularly alternating (predictable) 50% reinforcement schedules (Longenecker et al., 1952; Tyler et al., 1953; Capaldi et al., 1958; Habu and Ono, 1969) and that randomly delaying reinforcement on a proportion of trials also produces significant resistance to extinction (Crum et a., 1951). The present finding of a PREE in a tone foot shock fear conditioning protocol also indicates that the PREE is a general conditioning phenomenon and not confined simply to appetitive instrumental protocols (Lewis 1960, Haselgrove et al. 2004). Bayesian inferential theory can readily encompass this more general role for the PREE, whereas frustration theory (Amsel, 1958, 1992) is restricted to the omission of rewards in instrumental paradigms.

The immunohistochemical and electrophysiological results of Experiments 1 and 2 (Figures 7, 8, 13, 15, 19, and 21), showing increased p-ERK labelling and tone-evoked theta (8-10Hz) LFP activity in ventral hippocampus in the 50% reinforcement group, are the first evidence of hippocampal involvement in a fear conditioning PREE: previous studies implicating the hippocampus in the PREE have been confined to appetitive instrumental conditioning protocols (Rawlins et al. 1980, Rawlins et al. 1989; Gray, 1970; Gray et al., 1975). The observation of increased hippocampal activity in the random, high entropy, 50% reinforcement protocol is consistent with previous studies demonstrating increased hippocampal responses to unpredictability or uncertainty (Strange et al. 2005, Harrison et al. 2006, Herry et al. 2007, Vanni-Mercier et al. 2009; Schiffer et al., 2012; Payzan-LeNestour et al., 2013). The hippocampus, in addition to its role in spatial memory (O’Keefe and Dostrovsky, 1971), is also critical in coding and representing the memory of non-spatial event sequences (Terada et al., 2017), and it is this latter function which is presumably involved in representing the entropy changes which drive the PREE.

The higher frequency of hippocampal theta in the 50% compared to the 100% group in acquisition (Figure 13 and 14) is consistent with the findings of previous electrophysiological studies of the PREE employing food rewarded instrumental protocols (Gray, 1970; Gray et al., 1975). Gray (1970) reported a 6-7 Hz range of hippocampal LFP theta frequencies with continuous reinforcement compared to a 7.5-8.5 Hz range with partial reinforcement, observing that theta frequency at the end of acquisition was negatively correlated with resistance to extinction and concluding that “partial reinforcement training increases theta frequency and also resistance to extinction”. The 5.5-7.5Hz hippocampal theta in the 100% group (Figure 14) and 8-10Hz hippocampal theta in the 50% group (Figure 13) during acquisition in the current study are consistent with these previous findings. It is also notable that higher frequency oscillatory activity was also observed in the 100% group during the extinction session (Figures 17 and 20). This finding is consistent with a Bayesian inferential account, since after 100% reinforcement in acquisition, there will be an increase in entropy or uncertainty in the extinction session when reinforcement is omitted.

Increases in the frequency of hippocampal LFP theta have also been reported in association with enhanced spatial learning (Pan and McNaughton, 1997; Olvera-Cortez et al., 2012; López-Vázquez et al., 2014). Pan and McNaughton (1997) reported that hippocampal theta frequency was an inverse logarithmic function of training time on a Morris water maze task, and that significant learning impairments occurred when hippocampal LFP theta frequency dropped below 6.5 Hz. Olvera-Cortez et al., (2012) also using a Morris water maze task, and López-Vázquez et al., (2014), using a radial arm maze task, both reported that enhanced learning was associated with a greater proportion of high frequency (6.5-9.5 Hz) hippocampal theta compared to low frequency (4-6.5 Hz) theta in LFP recordings. The increased hippocampal theta frequency observed in the present study, both in the 50% group in both acquisition and extinction (Figures 13 and 19) in the 100% group in extinction (Figure 20) are consistent with these previous data, and indicate that increases in hippocampal theta frequency may represent a general mechanism for enhancement of hippocampal-dependent learning (Pan and McNaughton, 1997).

The learning-related increases in hippocampal theta frequency described above appear to be critically dependent on noradrenaline release from the locus coeruleus (Gray et al., 1975; Walling et al., 2011). Disruption of hippocampal noradrenaline release by lesions of the dorsal noradrenergic bundle or by noradrenergic blockers abolishes high frequency (7-9 Hz) hippocampal theta in awake rats (Gray et al., 1975), whereas stimulation of the locus coeruleus drives 7-9 Hz theta activity in hippocampus (Walling et al., 2011). Noradrenergic activation has also been linked to the facilitation of behavioural and attentional adaptation to changes in environmental variables (Sara, 2009), including “unexpected uncertainty” (Yu and Dayan, 2005; Hirsh et al., 2011; Payzan-LeNestour et al., 2013). These convergent findings that noradrenergic activation is associated not only with the PREE, but also with increased hippocampal theta frequency and with changes in environmental entropy suggest that uncertainty-dependent noradrenergic modulation of hippocampal theta frequency may represent a plausible neural mechanism for how random, unpredictable reinforcement sequences drive the PREE (Courville et al., 2006).

PREE-related increases in LFP theta power also occurred in BLA (Figures 10,12,16 and 18). This increased theta power in BLA in the 50% group suggests that, like hippocampus, amygdala is sensitive to changes in entropy or uncertainty, consistent with a previous report of uncertainty-dependent increases in amygdala activity (Herry et a., 2007). While increased theta synchronization of BLA activity following fear conditioning has been previously reported (Pare and Collins 2000), the current study provides the first evidence of theta oscillatory activity in BLA being modulated with a fear conditioning PREE. It is notable. however, that no significant differences in the density of c-fos or p-ERK labelling in BLA were observed between partial and continuous groups in the current study (Figure 6). The failure to observe a difference in the current study may reflect the multiple subpopulations and responses present in BLA and a lack of specificity in the immunolabelling technique. Electrophysiological studies of BLA units have reported that 17% are “fear” cells, responding to a fear conditioned CS+ in both acquisition and early extinction (Herry et al., 2008), 14% are “extinction” cells, responding to a fear conditioned CS+ only during late extinction (Herry et al., 2008) and 10% are “reward omission” cells, responding to food reward in acquisition and the omission of that reward in extinction Tye et al., (2010). The identification of a specific subpopulation of fear conditioned PREE cells in BLA may require more specific targeting with optogenetic (Kim et al., 2017) or DREADD (Designer Receptors Exclusively Activated by Designer Drugs) techniques (Roth, 2016).

In contrast to BLA, immunohistochemical labelling in central amygdala showed significant differences between partial and continuous groups (Figures 3, 4 and 5). The central amygdala is recognized as the principal output nucleus of the amygdala, controlling the expression of conditioned responses such as freezing (LeDoux, 2000), and therefore might be expected to show greater activity in the partial groups that exhibited increased resistance to extinction, i.e. increased conditioned responses (Figures 1 and 2). The greatest differential labelling was in the capsular subdivision of central amygdala, which receives convergent inputs of polymodal sensory information from BLA, nociceptive information from the parabrachial nucleus and efferents from infralimbic cortex (IL) (Bernard et al., 1993; Pinard et al., 2012). The projection from IL to amygdala has previously been implicated in mediating the extinction of conditioned fear (Milad and Quirk, 2002). Pinard et al., (2012) reported that the central amygdala region receiving the infralimbic projection, the Capsular Infralimbic Target Zone (CITZ), also shows co-localization of D1 dopamine receptors and they proposed that dopaminergic hyperpolarization of CITZ cells may result in blocking of infralimbic efferents and an attenuation of extinction. It is striking, therefore, that the PREE-related attenuation of extinction was associated with increased p-ERK labelling in the capsular subdivision of central amygdala (Figures 4 and 5). Future studies, using selective optogenetic targeting of cells in the CITZ and IL, may be able to directly test the hypothesis that inhibition of the IL input to the CITZ is a critical component of the neural circuitry underlying the PREE.

In addition to separate and distinct PREE-dependent responses in hippocampus and amygdala (Figures 7, 8, 21, 25, 28 and 29), the current data show significant PREE-related interactions between these regions (Figures 8, 22-25, 28,29). These interactions are consistent with the known functional anatomical connections between amygdala and hippocampus (Maren and Fanselow 1995; Pitkanen et al. 2000). The main output structure of the hippocampus, the ventral subiculum, has a direct anatomical connection with amygdala via the ventral angular bundle (VAB), with the greatest density of projections terminating in the lateral, basal and accessory basal amygdala nuclei (Pitkanen et al. 2000). The basal and accessory basal nuclei also send dense projections back to subiculum (Pitkanen et al. 2000). From these anatomical considerations, the subiculum appears well placed to coordinate its activity with that of amygdala. *In vivo* electrophysiological studies in rats support this hypothesis, showing that theta stimulation of the VAB produces long term potentiation (LTP) of neural responses in BLA (Maren and Fanselow 1995). Glutamatergic antagonists infused into BLA attenuate VAB-evoked responses, while electrolytic lesions of ventral subiculum prevent contextual fear conditioning (Maren and Fanselow 1995). The present results, showing significant hippocampal-amygdala interactions in a fear conditioned PREE, are consistent, therefore, with the established role of hippocampus in modulating amygdala-dependent conditioning (Maren and Fanselow, 1995).

Significant increases in the synchronization and coherence of hippocampus-BLA activity have been reported during fear memory retrieval (Seidenbecher et al., 2003; Narayanan et al., 2007a; Narayanan et al., 2007b; Popa et al., 2010). Seidenbecher et al., (2003), using a 100% reinforcement fear conditioning protocol in mice, reported that post-conditioning freezing behaviour to a CS tone was associated with synchronization of LFPs in BLA and CA1 of hippocampus in a low theta frequency band (mean = 4.3 Hz). The LFP coherence analyses of the current study, which show maximal values at low frequencies (2-5Hz) for the 100% group (Figures 22 and 24) are consistent with this previous report (Seidenbecher et al., 2003). Seidenbecher et al. (2003) also reported that higher frequency (8-14Hz) CS+-evoked oscillatory activity was observed in hippocampus, but not in BLA. In the current study, by contrast, higher frequency theta (8-10Hz) was observed in both hippocampus and BLA in the 50% group (Figures 10,13,16 and 19), and coherence between BLA and hippocampus peaked at these higher frequencies (Figures 22 and 24). Increased hippocampus-amygdala LFP coherence at 7-8.5Hz theta frequencies has been reported in rats in response to CS+ tones 24 hours following fear conditioning (Narayanan et al., 2007a) and after a reconsolidation/re-exposure procedure (Narayanan et al., 2007b). Popa et a., (2010) reported that hippocampus-amygdala theta coherence (6-8Hz) during overnight paradoxical sleep following tone foot shock fear conditioning in rats, correlated significantly with subsequent freezing to CS+ tones. The present data are consistent with these previous data, and go further in identifying increases in tone-evoked coherence in the 50% group during acquisition of conditioning (Figures 22 and 23). None of the studies described above included coherence analyses of acquisition data, although Narayanan (2007b) reported that no changes in hippocampus-BLA coherence were evident 2 hours after 100% conditioning. The current data suggest, therefore, that partial (uncertain) reinforcement not only induces higher frequency hippocampal theta oscillations but also an earlier occurrence of hippocampus-amygdala coherence than is observed with 100% reinforcement.

Granger Causality (Granger, 1969) has been used to analyze electrophysiological and neuroimaging data in order to assess directional causal influences between brain regions (Bernasconi and König, 1999; Guo et al., 2008; Luo et al., 2013). With regards to causal influences between hippocampus and amygdala, Popa et al., (2010) reported that, following fear conditioning in rats, the strength of the Granger Causality influence of BLA over hippocampal CA1 during overnight paradoxical sleep predicted the strength of conditioned freezing responses the next day, while Ghosh et al., (2013) reported increased bidirectional Granger Causality between CA1 and BLA after 1-2 days of chronic immobilization stress in rats. Our current data, showing increased bidirectional Granger Causality between subiculum and BLA on the day following fear conditioning acquisition are consistent with these previous reports, and go further in showing that this Granger Causality is enhanced by partial (uncertain) reinforcement in the 50% group (Figures 28 and 29). It is notable, however, that whereas increased tone-evoked coherence between subiculum and BLA was observed in both the acquisition (Figures 22 and 23) and extinction sessions in the 50% group (Figures 24 and 25), Granger Causality between these regions increased only in the extinction session on Day 2 (Figures 26-29). It is also notable that there were no significant differences in Granger Causality between pretone and tone periods (Figures 26-29) in contrast to coherence, which was significantly greater in the tone than the pretone periods (Figures 22-25). This finding suggests that the neural processes underlying Granger Causality are not only slower to develop, but are more stable and pervasive and not simply stimulus-driven.

The latency in the development of Granger Causality seen in both the current and previous studies (Figures 24 and 25; Popa et al., 2010; Ghosh et al., 2013) suggest that they may reflect slow-acting consolidation-related modifications of synaptic plasticity that are involved in stabilizing memory traces (Dudai, 2004). The more pervasive nature of Granger Causality, affecting both pretone and tone periods (Figures 26-29) would be consistent with this proposal. Changes in coherence, by contrast, may reflect fast-acting, stimulus-driven synchronizations of oscillatory activity that occur prior to the slower synaptic modifications occurring in memory consolidation. This proposal is consistent with the finding that long term potentiation (LTP), a widely held neurophysiological correlate of memory, is optimally induced by prior theta frequency bursts of spiking activity (Larsen and Munkácsy, 2014).

The critical role for uncertainty in producing the PREE, as proposed by Bayesian inferential theory and supported both by previous studies (Longenecker et al., 1952; Tyler et al., 1953; Capaldi et al., 1958; Habu and Ono, 1969; Crum et a., 1951) and by the current data, suggests that a better name for the phenomenon might be the “Unpredictable Reinforcement Extinction Effect”. The current findings suggest, therefore, that explicitly manipulating stimulus entropy rather than the proportion of unreinforced to reinforced trials may prove very fruitful in investigating the detailed neural mechanisms of the PREE and of learning in general.

## Conclusion

The present study has a number of notable findings: it is the first report of a PREE in tone foot shock fear conditioning in rats (Figures 1 and 2); by demonstrating a PREE in a Pavlovian protocol, it supports the view that the PREE is a general phenomenon in reinforcement learning, and is not restricted to appetitive instrumental conditioning protocols (Lewis 1960, Haselgrove et al. 2004); it provides immunohistochemical and electrophysiological evidence for hippocampal involvement in mediating the PREE, consistent with precious appetitive instrumental studies (Figures 7, 8, 21, 25, 28 and 29; Rawlins et al. 1980, Rawlins et al. 1989; Gray, 1970; Gray et al., 1972; Gray et al., 1975); more specifically, it indicates a critical role for higher frequency (8-10 Hz) hippocampal theta oscillations in mediating the PREE, consistent with previous studies (Gray, 1970; Gray et al., 1972; Gray et al., 1975); it provides evidence that the PREE is associated with increased coherence of hippocampal-amygdala activity that is maximal at theta frequencies and that occurs early in acquisition; it demonstrates bidirectional Granger Causality between hippocampus and amygdala activity, which is only evident 24 hours after acquisition of fear conditioning, and which is significantly enhanced by the 50% reinforcement protocol (Figures 26-29); finally, it shows an entropy-like inverted-U distribution of both conditioned responses (Figure 2) and hippocampal/amygdala activity (Figures 4 and 7) against reinforcement probability that supports a Bayesian interpretation of the PREE as a phenomenon driven principally by the uncertainty of stimulus contingencies (Courville et al., 2006).

## Acknowledgements

This work was supported by the award of a scholarship from the National Health and Medical Research Council of Australia. The authors would like to thank the all our colleagues at the Queensland Brain Institute for their support and encouragement with this project.

